# Myeloid DC-SIGN drives tumor immune evasion and resistance to PD-1 blockade through ICAM-2–ERM–mediated T-cell mechanosuppression

**DOI:** 10.64898/2026.01.19.699861

**Authors:** Ting Pan, Tingting Wang, Jiaqi Wu, Heping Wang, Bo Zeng, Yangyang Li, Ming Yi, Ruirui He, Lingyun Feng, Zhihui Cui, Guoling Huang, Panyin Shu, Xi Li, Yuan Wang, Yanyun Du, Zhen Li, Xue Xiao, Chenhui Wang

## Abstract

Resistance to PD-1–based immunotherapy remains a major challenge in cancer treatment, especially in immunologically “cold” tumors characterized by T-cell exclusion and impaired effector function. Tumor-associated myeloid cells are important mediators of this resistance, but whether they directly suppress T-cell immunity through mechanical regulation remains unclear. Here, we identify DC-SIGN (CD209), a myeloid-expressed C-type lectin receptor, as a mechanosuppressive regulator of antitumor T-cell immunity. DC-SIGN is selectively enriched in tumor-infiltrating M2-like macrophages and monocytes, and its expression is associated with poor clinical response to PD-1 blockade and elevated TIDE scores across multiple cancer types. Mechanistically, DC-SIGN directly engages ICAM-2 on T cells and activates ROCK1–ERM signaling. This interaction increases T-cell cortical stiffness, destabilizes T cell–antigen-presenting cell interactions, and attenuates T-cell receptor signaling. Blocking the DC-SIGN–ICAM-2–ERM pathway restores T-cell activation and promotes CD8+ T-cell infiltration and effector function in mouse, humanized, and *ex vivo* human tumor systems. Importantly, DC-SIGN blockade overcomes resistance to PD-1–based immunotherapy in refractory and immunologically “cold” tumor models, and further enhances the response to PD-1 inhibition. Together, our findings define a myeloid-derived mechanical immune checkpoint and reveal T-cell mechanosuppression as a targetable mechanism of tumor immune evasion and immunotherapy resistance.

## Introduction

Immune checkpoint blockade targeting the PD-1/PD-L1 axis has transformed cancer therapy. However, most patients with solid tumors still fail to achieve durable responses, particularly those with immunologically “cold” tumors marked by limited T-cell infiltration and impaired effector function(1–3). These limitations suggest that suppressive mechanisms beyond canonical PD-1/PD-L1 signaling continue to constrain productive antitumor T-cell immunity. Tumor-associated myeloid cells are major contributors to this resistance. They restrict T-cell infiltration, activation, and function through inhibitory ligands, cytokines, metabolic competition, and stromal remodeling(4–7). However, how myeloid cells directly impose suppressive constraints on T cells, especially in PD-1–refractory and T-cell-excluded tumors, remains incompletely understood.

Most established immune checkpoints are viewed as biochemical receptor–ligand pathways. These pathways attenuate intracellular signaling in T cells and thereby limit TCR signaling, cytokine production, proliferation, and effector differentiation(8–15). By contrast, it remains largely unclear whether tumor-associated myeloid cells can suppress T-cell activation through contact-dependent mechanical regulation. Productive T-cell activation requires not only antigen recognition and costimulatory signaling, but also stable engagement with antigen-presenting cells, sustained immunological synapse formation, and dynamic remodeling of the actin cortex(16–19). These mechanical processes control T-cell spreading, receptor organization, signaling-zone stability, and the duration of T cell–APC interactions. Thus, perturbations in T-cell cortical mechanics may represent a distinct mode of immune suppression. This mode may operate in parallel with, or independently of, canonical inhibitory checkpoint signaling. Defining such mechanisms may reveal previously unrecognized myeloid-derived checkpoints and provide therapeutic opportunities for tumors that remain resistant to PD-1–based immunotherapy.

DC-SIGN, also known as CD209, is a C-type lectin receptor expressed by dendritic cells, monocytes, and macrophages(20,21). It regulates cell–cell adhesion and immune-cell trafficking through interactions with ICAM family members, including ICAM-3 and ICAM-2, and serves as an attachment receptor for diverse pathogens(22–28). In cancer, DC-SIGN⁺ myeloid cells are associated with immunosuppressive tumor microenvironments, impaired antitumor immunity, poor clinical outcomes, and reduced responsiveness to immunotherapy(29–34). Previous studies have linked DC-SIGN⁺ tumor-associated macrophages to CD8⁺ T-cell dysfunction. Tumor-derived ligands, including CEA, CEACAM1, and Mac-2BP, can also engage DC-SIGN to suppress dendritic-cell function(30–32). However, whether DC-SIGN directly acts on T cells, and whether its adhesive interactions with ICAM family members impose mechanical constraints on T-cell activation, remains unknown.

Here, we identify DC-SIGN as a myeloid-derived mechanosuppressive regulator that directly restrains antitumor T-cell immunity. DC-SIGN engages ICAM-2 on T cells and activates ROCK1–ERM signaling. This pathway increases T-cell cortical stiffness, destabilizes T cell–antigen-presenting cell interactions, and attenuates T-cell receptor signaling. Disruption of the DC-SIGN–ICAM-2–ERM axis restores T-cell activation, enhances CD8⁺ T-cell infiltration and effector function, and promotes antitumor immunity across mouse, humanized, and ex vivo human tumor systems. DC-SIGN blockade also overcomes resistance to PD-1–based immunotherapy in refractory and immunologically “cold” tumors, including ovarian cancer and glioblastoma. Analyses of human tumors show that elevated DC-SIGN expression is associated with CD8⁺ T-cell exclusion, poor clinical outcome, and resistance to PD-1 blockade. Together, these findings define a myeloid-driven mechanical immune checkpoint and establish T-cell cortical mechanics as a targetable mechanism of tumor immune evasion.

## Results

### Myeloid CD209/DC-SIGN is associated with immunosuppressive tumor microenvironments and resistance to PD-1 blockade

To identify myeloid-derived mechanisms associated with resistance to PD-1 blockade, we analyzed single-cell transcriptomes of surgically resected NSCLC tumors from 25 patients treated with neoadjuvant chemotherapy plus anti–PD-1 therapy (GSE243013) (35) (Fig. 1A). Among these patients, 9 achieved a major pathological response (MPR), whereas 16 did not. Because tumor-associated myeloid cells are key regulators of immune evasion, we focused on the myeloid compartment and compared macrophages from non-MPR and MPR tumors. This analysis identified a set of genes preferentially upregulated in macrophages from non-MPR tumors (Fig. 1B). Among the genes upregulated in macrophages from non-MPR tumors, several macrophage-enriched molecules, including MERTK, FOLR2, CCL8, ENPP2, OLFML3, and PDGFB, have previously been reported to participate in TAM-mediated immunosuppressive tumor microenvironments through promoting suppressive cytokine production, T-cell dysfunction, stromal remodeling, angiogenesis, immune evasion, and resistance to immune checkpoint blockade(11,36–41). Intersecting the genes upregulated in macrophages with a curated human membrane-associated gene set yielded six candidate surface molecules (Fig. 1C). Further prioritization using TIDE-based T-cell dysfunction risk scoring identified CD209 as the top-ranked candidate, nominating it as a potential myeloid-associated checkpoint linked to anti–PD-1 resistance (Fig. 1D).

**Figure 1.**
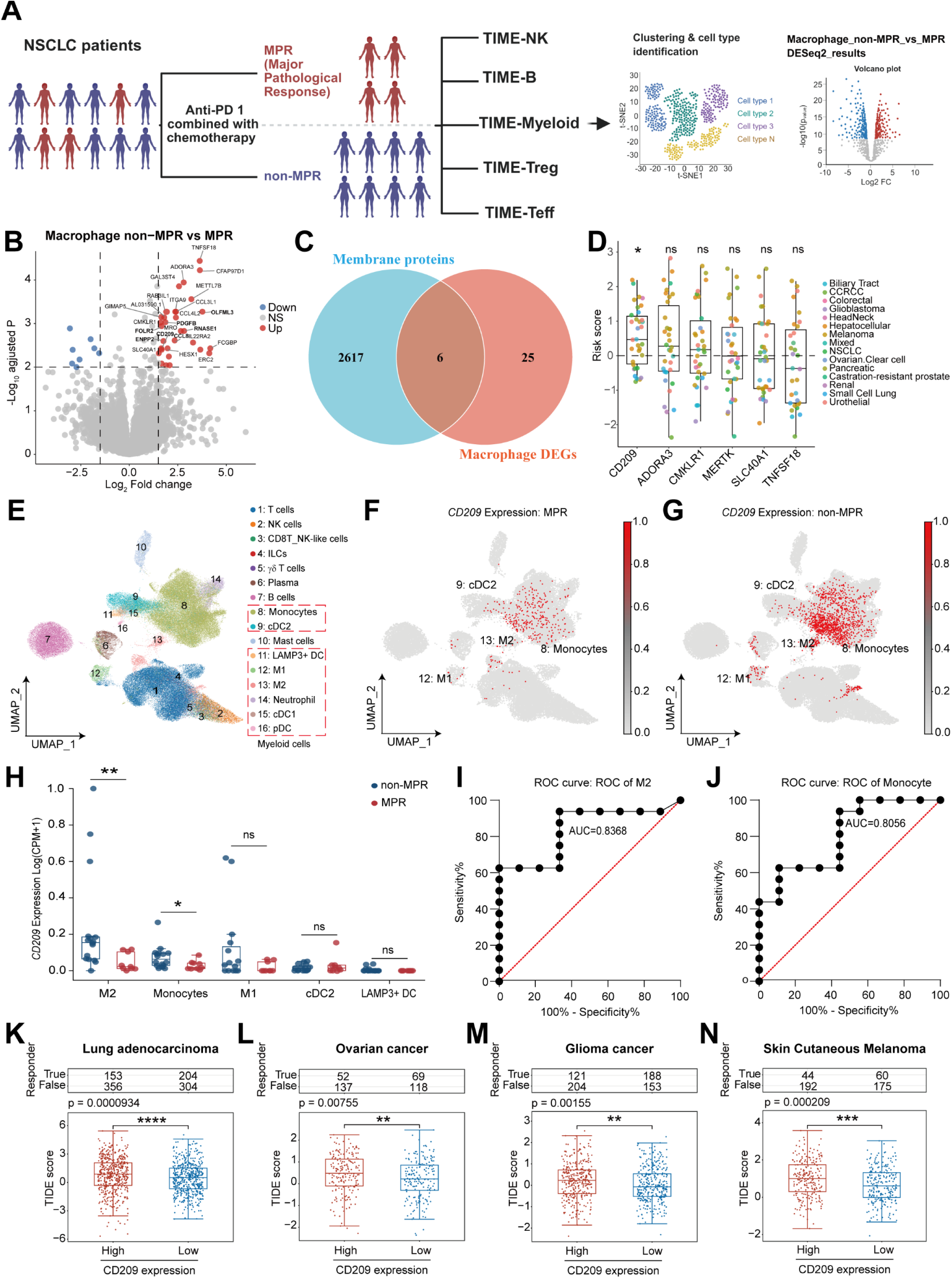
Myeloid CD209 is associated with immunosuppressive tumor microenvironments and resistance to PD-1 blockade. **(A)** Schematic overview of the single-cell transcriptomic analysis workflow. Single-cell RNA-sequencing data from NSCLC tumors of patients receiving neoadjuvant anti–PD-1 combined with chemotherapy were stratified into major pathological response (MPR) and non-MPR groups, followed by cell clustering and differential expression analysis in macrophages. **(B)** Volcano plot showing differentially expressed genes (DEGs) in macrophages from non-MPR versus MPR tumors. **(C)** Venn diagram showing the overlap between macrophage DEGs and membrane protein–encoding genes, identifying six candidate membrane molecules. **(D)** T-cell dysfunction risk z-scores of candidate genes calculated using the TIDE algorithm. P values were calculated using a two-sided Wilcoxon signed-rank test against zero, followed by Benjamini–Hochberg correction for multiple testing. **(E)** UMAP visualization of major immune cell populations identified from NSCLC single-cell transcriptomic data, highlighting myeloid cell subsets. **(F–G)** Feature plots showing CD209 expression in tumors from MPR (F) and non-MPR (G) patients. **(H)** Quantification of CD209 expression levels across indicated myeloid cell subsets in MPR and non-MPR tumors. **(I–J)** Receiver operating characteristic (ROC) curve analysis evaluating the predictive performance of CD209 expression in M2 macrophages (I) and monocytes (J) for anti–PD-1 therapeutic response. **(K–N)** Association between CD209 expression and TIDE scores in lung adenocarcinoma (K), ovarian cancer (L), glioma (M), and skin cutaneous melanoma (N). Patients were stratified into CD209-high and CD209-low groups. Statistical significance was determined using two-sided Wilcoxon single-rank tests (D). Data are presented as box plots showing median and interquartile range (D). *P < 0.05, **P < 0.01, ***P < 0.001, ****P < 0.0001; ns, not significant. Statistical analyses were performed using two-tailed unpaired t tests (H).

We next examined the cellular distribution of CD209 within the same NSCLC cohort. CD209 expression was largely restricted to myeloid cells and was highest in M2-like macrophages (Fig. 1E–H). Notably, CD209 expression in M2-like macrophages and monocytes was significantly elevated in non-MPR tumors compared with MPR tumors, whereas no significant difference was observed in dendritic cells or M1-like macrophages (Fig. 1H). Receiver operating characteristic analysis further showed that CD209 expression in M2-like macrophages and monocytes discriminated nonresponders from responders to PD-1–based therapy (Fig. 1I–J), supporting its potential association with therapeutic resistance. Consistently, CD209⁺ cell abundance was significantly higher in an independent cohort of PD-1–refractory lung cancer patients than in responders (Fig. S1A–C).

To further define the cellular distribution of CD209 within the tumor microenvironment, we analyzed single-cell RNA-sequencing data generated from paired tumor and adjacent normal tissues in our laboratory. Unsupervised clustering identified 18 major immune cell populations based on established lineage markers. CD209 expression was predominantly enriched in immunosuppressive M2-like macrophages and monocytes, but was largely absent from M1-like macrophages (Fig. S1D–F). Compared with adjacent tissues, tumors showed significantly increased CD209 expression in M2-like macrophages and monocytes, whereas CD209 expression in dendritic cells was reduced (Fig. S1F). This pattern is consistent with previous studies linking CD209⁺ myeloid cells to tumor immune evasion and poor clinical outcomes (21,29,31,32). Analysis of an independent LUAD single-cell RNA-sequencing dataset (GSE194070) further confirmed the preferential enrichment of CD209 in tumor-infiltrating M2-like macrophages and monocytes (42) (Fig. S1G–I). Together, these data indicate that CD209 is selectively upregulated in immunosuppressive myeloid populations within the tumor microenvironment, supporting its potential involvement in myeloid-mediated immune suppression.

To evaluate the broader clinical relevance of CD209, we performed pan-cancer analyses and found that elevated CD209 expression correlated with increased TIDE scores across multiple tumor types, including lung adenocarcinoma (LUAD), ovarian cancer, glioma, melanoma, prostate adenocarcinoma (PRAD), breast invasive carcinoma (BRCA), head and neck squamous cell carcinoma (HNSC) and uterine corpus endometrial carcinoma (UCEC) (Fig. 1K–N and Fig. S1J–N). Together, these findings nominate CD209 as a myeloid-associated marker of dysfunctional antitumor immunity and resistance to PD-1–based immunotherapy, raising the possibility that CD209 may functionally contribute to therapeutic resistance and serve as a candidate biomarker for patient stratification.

### CD209 blockade restores T-cell activation and overcomes resistance to PD-1 blockade

To determine whether CD209 contributes to resistance to immune checkpoint blockade, we first assessed its direct effects on T-cell activation. Recombinant human CD209-Fc bound efficiently to Jurkat T cells as well as primary murine CD4⁺ and CD8⁺ T cells (Fig. 2A–B), suggesting the presence of conserved receptors across species. Functionally, plate-bound CD209-Fc markedly inhibited T-cell activation following anti-CD3/CD28 stimulation in Jurkat cells, human peripheral blood mononuclear cells (PBMCs), and primary mouse CD4⁺ and CD8⁺ T cells (Fig. 2C). These findings establish CD209 as a direct inhibitor of T-cell activation and indicate that this inhibitory mechanism is evolutionarily conserved. Consistent with previous reports implicating CD209⁺ myeloid cells in suppressing T-cell responses(43), our data provide direct evidence that CD209 can function as a T-cell inhibitory regulator.

**Figure 2.**
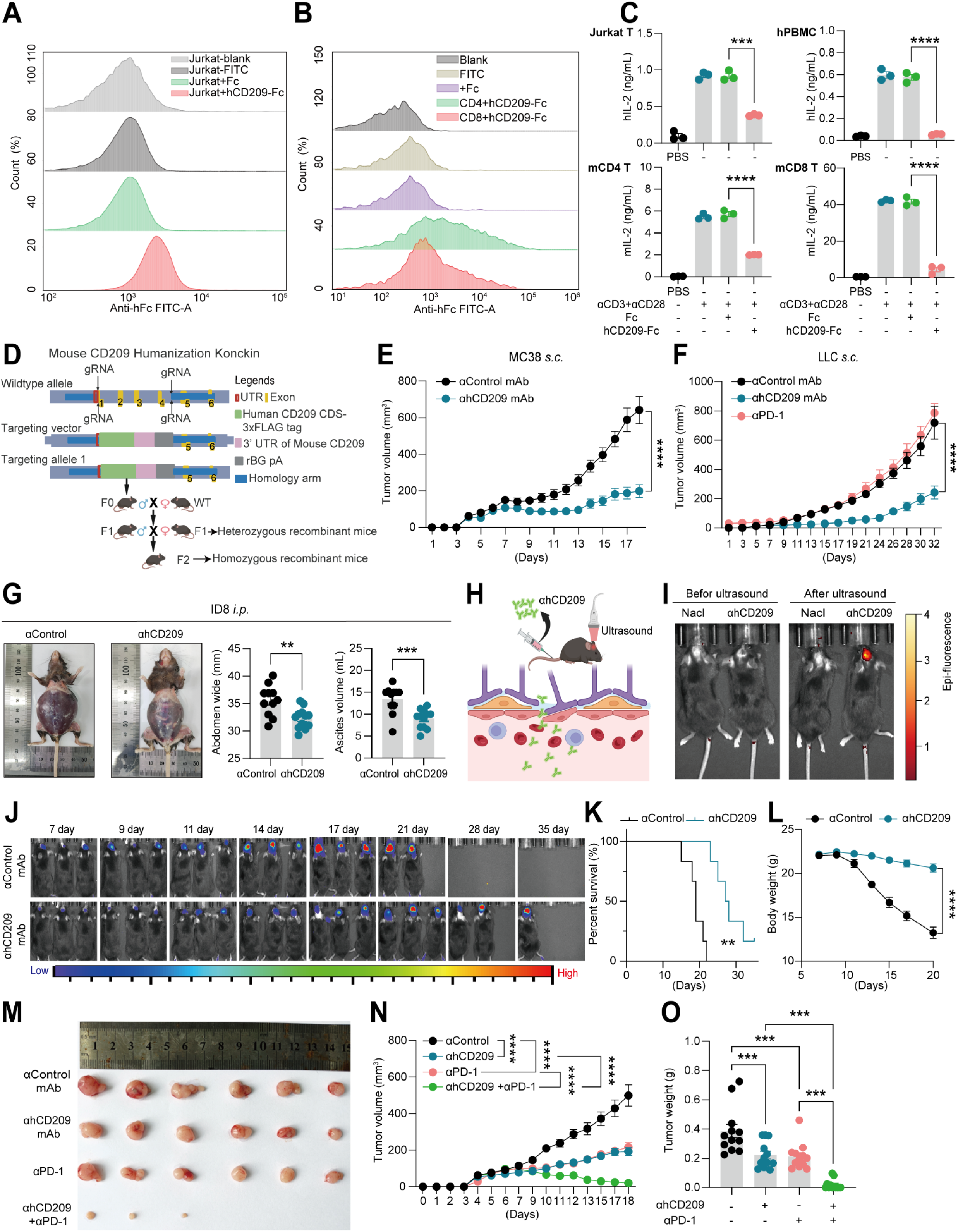
CD209 blockade restores T-cell activation and overcomes resistance to PD-1 blockade. **(A)** Jurkat T cells were incubated with Fc or hCD209-Fc, followed by flow cytometric analysis of cell-bound CD209-Fc. **(B)** Mouse CD4⁺ and CD8⁺ T cells were incubated with Fc or hCD209-Fc, followed by flow cytometric analysis of CD209-Fc binding. **(C)** Jurkat cells, human PBMCs, mouse CD4⁺ T cells, or CD8⁺ T cells were left untreated or stimulated with plate-bound αCD3+αCD28 in the presence of Fc or hCD209-Fc for 48 hours. IL-2 production was measured by ELISA. **(D)** Schematic of the strategy used to generate hCD209 knock-in (KI) mice. Homozygous KI mice were used for tumor experiments. **(E)** Tumor growth curves of MC38 subcutaneous tumors in hCD209-KI mice treated with control or anti-hCD209 monoclonal antibody (mAb) (n=13/12). **(F)** Tumor growth curves of LLC subcutaneous tumors in hCD209-KI mice treated with control, anti-PD-1, or anti-hCD209 mAb (n=13/6/13). **(G)** ID8 ovarian tumor cells were injected intraperitoneally into hCD209-KI mice followed by control or anti-hCD209 mAb treatment. Representative abdominal images, abdominal width, and ascites volume at day 30 are shown (n=11/12). **(H)** Schematic of focused ultrasound–mediated transient disruption of the blood–brain barrier. **(I)** In vivo fluorescence imaging of FITC-labeled anti-hCD209 mAb distribution in hCD209-KI mice before or after ultrasound-mediated blood–brain barrier disruption. **(J–L)** GL261-Luc glioma cells were implanted intracranially into hCD209-KI mice, followed by control or anti-hCD209 mAb treatment after ultrasound. Tumor progression (J), survival (K), and body weight (L) are shown (n=6/6). **(M–O)** MC38 tumor-bearing hCD209-KI mice were treated with control, anti-hCD209 mAb, anti–PD-1 mAb, or combination therapy. Representative tumor images (M), tumor growth curves (N), and tumor weights (O) are shown (n=12/12/12/12). Statistical significance was determined by two-way ANOVA (E, F, L, N), one-way ANOVA (O), two-tailed unpaired t test (C, G), and log-rank (Mantel–Cox) test (K). Data in (A–O) are pooled from two independent experiments. *P < 0.05; **P < 0.01; ***P < 0.005; ****P < 0.001. All error bars represent mean ± SEM.

To determine whether targeting CD209 has therapeutic potential, we generated a neutralizing monoclonal antibody against human CD209 that effectively reversed CD209-mediated suppression of T-cell activation in both human and mouse systems (Fig. S2A–D). We next established a human CD209 knock-in mouse model (hCD209-KI), in which human CD209 is expressed under the endogenous mouse *cd209* promoter (Fig. 2D). Therapeutically, CD209 blockade significantly inhibited tumor growth across multiple models. In MC38 tumor–bearing hCD209-KI mice, anti-CD209 treatment markedly suppressed tumor progression compared with control (Fig. 2E). Notably, in the PD-1–refractory “cold” LLC model, CD209 blockade remained highly effective, resulting in pronounced tumor growth inhibition (Fig. 2F). In addition, CD209 blockade significantly reduced metastatic burden in B16 hepatic and pulmonary metastasis models (Fig. S2E). Collectively, these results demonstrate that targeting CD209 elicits potent antitumor activity *in vivo*, including in immunotherapy-resistant tumor settings.

A major limitation of immune checkpoint blockade (ICB) is its poor efficacy in immunologically “cold” tumors such as ovarian cancer and glioblastoma(44,45). We therefore evaluated whether CD209 blockade is effective in these settings. In a murine ovarian cancer model, anti-CD209 treatment markedly reduced ascites accumulation and abdominal distension compared with isotype controls (Fig. 2G). We next assessed the therapeutic potential of CD209 blockade in an orthotopic glioblastoma model. Because antibodies do not efficiently cross the blood–brain barrier (BBB), we employed focused ultrasound to transiently open the BBB and facilitate antibody delivery(46,47). Fluorescence imaging confirmed efficient intracerebral accumulation of FITC-labeled anti-CD209 antibodies following ultrasound treatment (Fig. 2H–I). In this model, CD209 blockade significantly prolonged survival and reduced weight loss compared with control treatment (Fig. 2J–L). Notably, CD209 blockade alone outperformed temozolomide (TMZ) monotherapy, and combination treatment further enhanced therapeutic efficacy (Fig. S2F–G). Furthermore, combined blockade of CD209 and PD-1 resulted in near-complete suppression of tumor growth in the MC38 model, indicating a strong synergistic antitumor effect (Fig. 2M–O). Collectively, these findings demonstrate that CD209 blockade overcomes resistance to PD-1–based immunotherapy, converts immunologically “cold” tumors into treatment-responsive states, and further enhances antitumor efficacy in combination with PD-1 blockade.

### CD209 blockade restores CD8⁺ T cell–mediated antitumor immunity by relieving myeloid-mediated suppression

To investigate the cellular mechanisms underlying the therapeutic effects of CD209 blockade, we performed single-cell RNA sequencing of MC38 tumors from hCD209-KI mice. Unsupervised clustering identified 12 major cell populations, with a significant increase in the relative abundance of T cells following CD209 blockade (Fig. 3A–C, Fig. S3A, S3D). Re-clustering of the T-cell compartment revealed eight transcriptionally distinct subsets, among which CD8⁺ effector T cells were markedly expanded (Fig. 3D–F, Fig. S3B, S3E), suggesting a central role for this population in mediating the antitumor response. Analysis of the myeloid compartment identified seven macrophage subsets, and CD209 blockade selectively reduced the abundance of M2-like macrophages without affecting other populations (Fig. 3G–I, Fig. S3C, S3F). Collectively, these findings show that CD209 blockade relieves myeloid-mediated suppression and promotes CD8⁺ T-cell–driven antitumor immunity.

**Figure 3.**
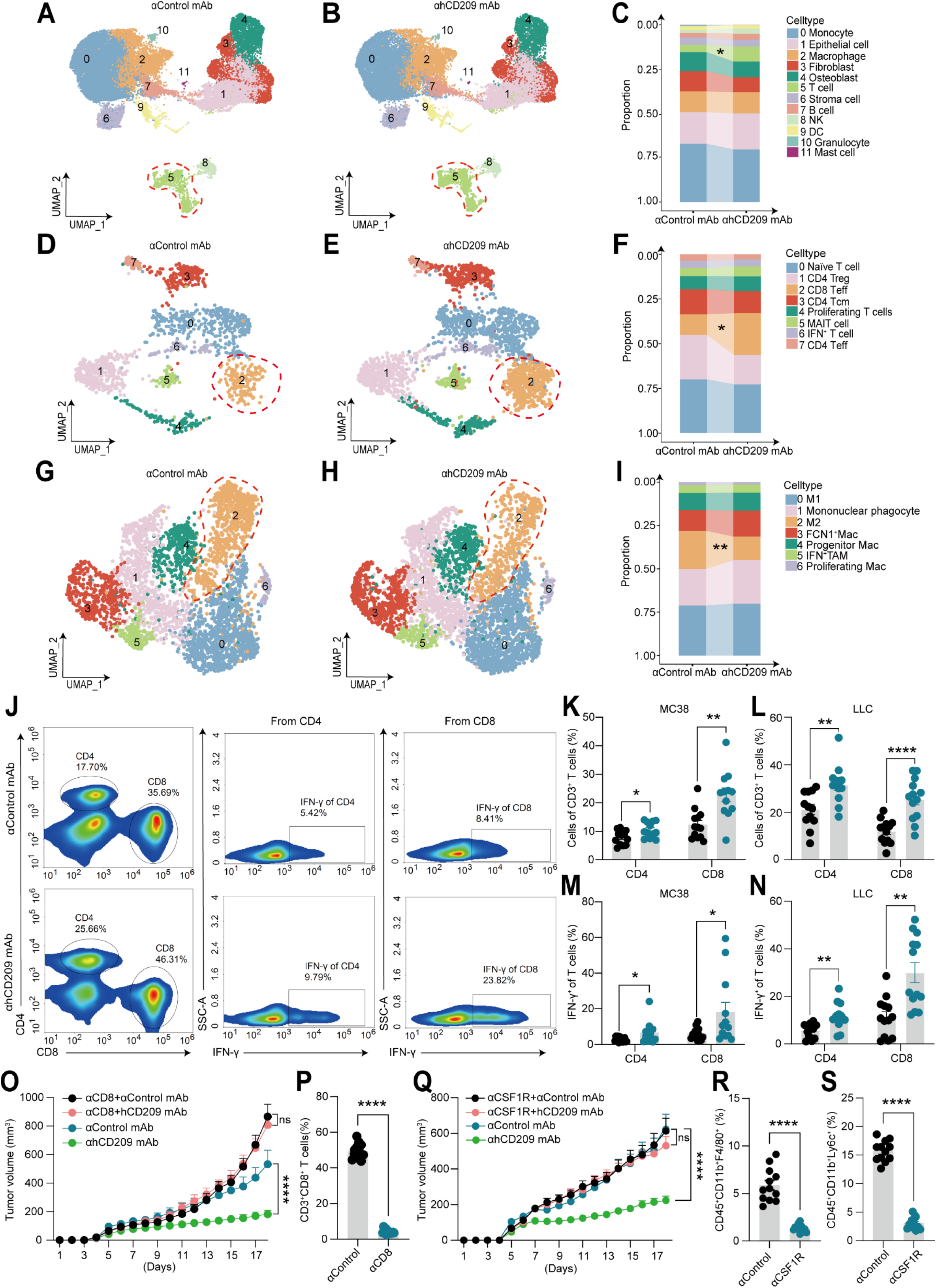
CD209 blockade restores CD8⁺ T cell–mediated antitumor immunity by relieving myeloid-mediated suppression. MC38 tumors were subcutaneously implanted into hCD209-KI mice, followed by treatment with control or anti-hCD209 monoclonal antibody (mAb). Single-cell RNA sequencing (scRNA-seq) was performed on paired tumors (n=3 per group). **(A–C)** UMAP visualization of all cell populations (A–B) and corresponding cell-type proportions (C) in control and anti-hCD209 treated tumors. **(D–F)** UMAP projection of T cell subsets (D–E) and their relative proportions (F) following control or anti-hCD209 treatment. **(G–I)** UMAP projection of macrophage subsets (G–H) and their relative proportions (I) following control or anti-hCD209 treatment. **(J–L)** Flow cytometric analysis of tumor-infiltrating CD4⁺ and CD8⁺ T cells (J) and their quantification in MC38 (K) and LLC (L) tumor models following control or anti-hCD209 mAb treatment. **(M–N)** Flow cytometric analysis of IFN-γ–producing CD4⁺ and CD8⁺ T cells in MC38 (M) and LLC (N) tumors following control or anti-hCD209 treatment. **(O–P)** hCD209-KI MC38 tumor–bearing mice were treated with control or anti-hCD209 monoclonal antibody (mAb), with or without CD8⁺ T cell depletion. Tumor growth curves (O) and depletion efficiency of CD8⁺ T cells (P) are shown (n=12/12/12/12). **(Q–S)** hCD209-KI MC38 tumor–bearing mice were treated with control or anti-hCD209 monoclonal antibody (mAb), with or without anti-CSF1R treatment. Tumor growth curves (Q) and depletion efficiency of CSF1R⁺ myeloid cells (R–S) are shown (n=12/12/12/12). Statistical significance was determined using two-tailed unpaired t tests (C, F, I, K-N, P, R-S) and two-way ANOVA (O, Q). *P < 0.05; **P < 0.01; ***P < 0.005; ****P < 0.001. All error bars represent mean ± SEM.

CD209 blockade significantly increased intratumoral CD4⁺ and CD8⁺ T-cell infiltration across tumor models, as determined by flow cytometry (Fig. 3J–L, Fig. S3G–I). This expansion was accompanied by a marked increase in IFN-γ–producing T-cell subsets and elevated systemic IFN-γ levels in serum and tumor culture supernatants (Fig. 3M–N, Fig. S3J–N), indicating enhanced effector function. Consistent with our single-cell RNA-seq data and human LUAD datasets, CD209⁺ monocytes and macrophages were enriched in tumors compared with spleens, whereas dendritic cells displayed reduced expression (Fig. S4A–B, Fig. S1F, Fig. S1I), supporting a conserved tumor-specific myeloid phenotype. Flow cytometric analysis of intratumoral myeloid subsets revealed that CD209 blockade selectively reduced polymorphonuclear (PMN)-MDSCs across tumor models, while leaving classical and non-classical monocyte populations largely unaffected (Fig. S4C–D). In addition, dendritic cell frequencies remained unchanged following CD209 blockade (Fig. S4F–G).

To determine whether these immune populations are required for therapeutic efficacy, we depleted CD8⁺ T cells (anti-CD8α) or monocytes/macrophages (anti-CSF1R) in tumor-bearing hCD209-KI mice. Notably, depletion of either compartment completely abrogated the antitumor effects of CD209 blockade (Fig. 3O–S), demonstrating that both CD8⁺ T cells and tumor-associated myeloid cells are essential mediators of the therapeutic response. Collectively, these findings suggest that CD209 directly suppresses CD8⁺ T-cell function within the tumor microenvironment, prompting us to investigate the underlying receptor-mediated mechanism.

### ICAM-2 is the functional receptor mediating CD209-dependent T-cell suppression

To identify candidate receptors mediating CD209-dependent T-cell inhibition, we performed immunoprecipitation–mass spectrometry using recombinant CD209-Fc as bait in Jurkat T-cell membrane lysates. This analysis identified multiple transmembrane proteins interacting with CD209, among which ICAM-3—previously reported to bind CD209 and regulate T cell–dendritic cell interactions—emerged as a candidate (Fig. 4A) (48). However, genetic deletion of ICAM-3 in Jurkat cells did not impair the ability of CD209 to suppress TCR-mediated activation (Fig. S5A–B), demonstrating that ICAM-3 is not required for CD209-mediated inhibition.

**Figure 4.**
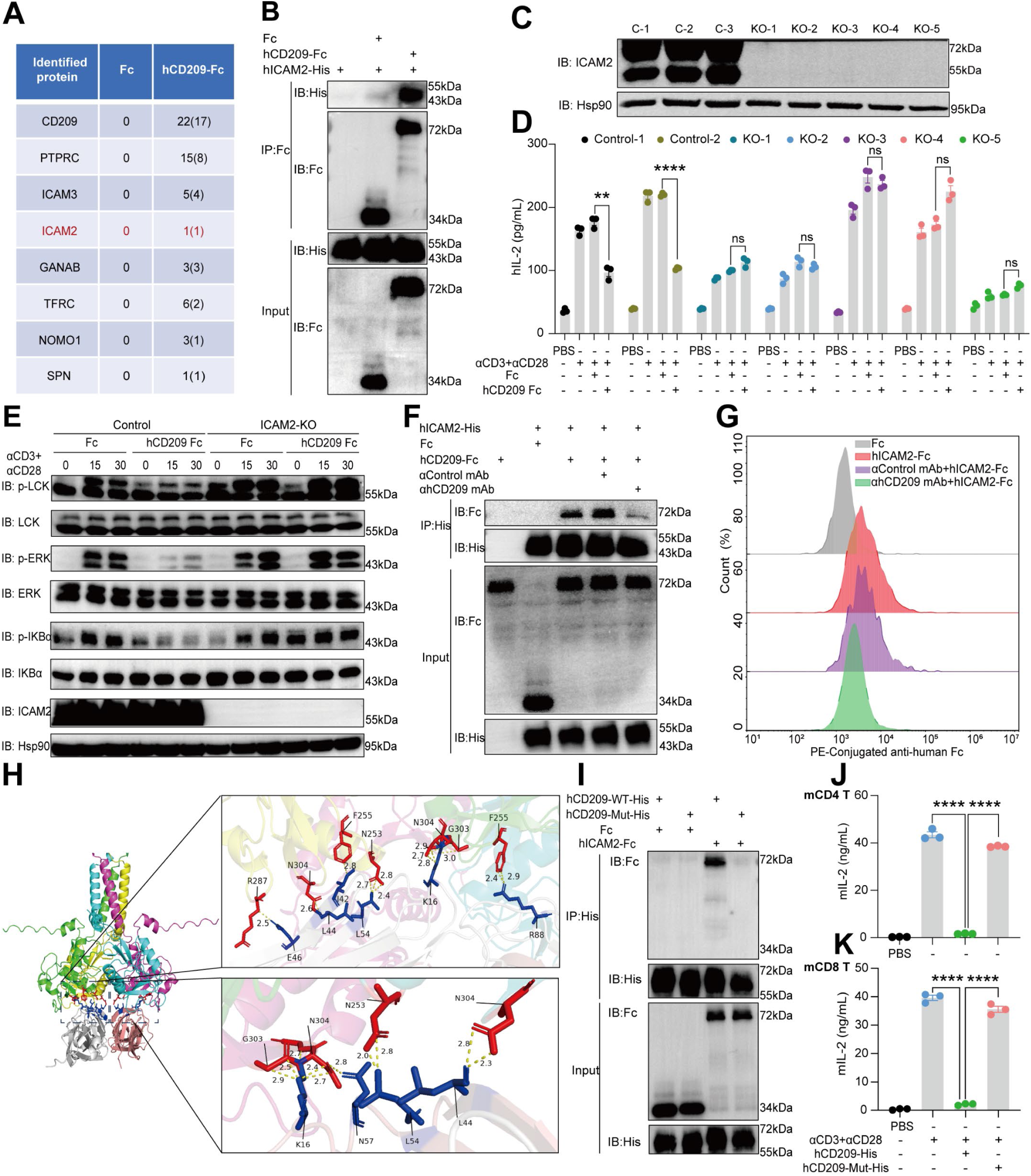
ICAM-2 is the functional receptor mediating CD209-dependent T-cell suppression. **(A)** Immunoprecipitation coupled with mass spectrometry (IP–MS) using membrane lysates from Jurkat cells identifies proteins interacting with CD209. **(B)** Immunoprecipitation assay using protein G beads in the presence of recombinant hCD209-Fc and hICAM2-His proteins, showing binding of hICAM2-His to CD209-Fc. **(C)** Immunoblot analysis confirming ICAM2 expression in control and ICAM2-knockout (KO) Jurkat cells. **(D)** Control or ICAM2-KO Jurkat cells were stimulated with plate-bound αCD3+αCD28 in the presence of Fc or hCD209-Fc, followed by ELISA measurement of IL-2 production. **(E)** Control or ICAM2-KO Jurkat cells treated with Fc or hCD209-Fc were stimulated with plate-bound αCD3+αCD28 for the indicated times, followed by immunoblot analysis of TCR signaling–related proteins. **(F)** Immunoprecipitation using Ni beads in the presence of recombinant hCD209-Fc and hICAM2-His proteins with control or anti-hCD209 mAb, demonstrating antibody-dependent disruption of CD209–ICAM2 interaction. **(G)** Flow cytometric analysis of ICAM2-Fc binding to 293T cells overexpressing human CD209 in the presence of control or anti-hCD209 mAb. **(H)** Structural model of the CD209–ICAM2 interaction predicted by AlphaFold3, highlighting key interacting residues (CD209, red; ICAM2, blue). **(I)** Immunoprecipitation using Ni beads with recombinant hCD209-WT-His or mutant His-tagged proteins and hICAM2-Fc, demonstrating reduced binding of the CD209 mutant to ICAM2 **(J–K)** Mouse CD4⁺ (J) or CD8⁺ (K) T cells were stimulated with αCD3+αCD28 in the presence of control, hCD209-WT-His, or mutant proteins for 48 hours, followed by ELISA measurement of IL-2 production. Statistical significance was determined using two-tailed unpaired t tests (D, J, K). *P < 0.05; **P < 0.01; ***P < 0.005; ****P < 0.001. All error bars represent mean ± SEM. Data in (B-G, I–K) are pooled from two independent experiments.

To identify the functional receptor responsible for CD209-mediated inhibition, we examined ICAM-2, a previously reported CD209 ligand(49), which was also recovered in our IP–MS screen (Fig. 4A). Direct interaction between hCD209 and hICAM-2 was confirmed by reciprocal co-immunoprecipitation (Fig. 4B). Strikingly, CRISPR-mediated deletion of ICAM-2 in Jurkat cells completely abrogated CD209-mediated suppression, as evidenced by full restoration of TCR-induced phosphorylation of LCK, ERK, and IκBα, as well as IL-2 production (Fig. 4C–E). These findings establish ICAM-2 as the essential receptor through which CD209 inhibits T-cell activation.

Consistently, neutralizing anti-CD209 antibodies disrupted the CD209–ICAM-2 interaction in both human and murine systems, effectively blocking the binding of human CD209 to both human and mouse ICAM-2 and thereby preventing CD209-mediated suppression of T-cell activation (Fig. 4F–G, Fig. S5C). These results support the cross-species compatibility of the CD209–ICAM-2 interaction and validate the physiological relevance of the hCD209 knock-in mouse model.

Structurally, CD209 forms tetramers via its neck domain(50), whereas ICAM-2 exists predominantly as a dimer. AlphaFold3 modeling of the CD209–ICAM-2 complex identified a core interaction interface involving five CD209 residues (Fig. 4H). Alanine substitution of these residues (CD209-5A) abolished ICAM-2 binding and eliminated CD209-mediated suppression of CD4⁺ and CD8⁺ T-cell activation, confirming that this interface is functionally required (Fig. 4I–K). Collectively, these findings establish ICAM-2 as the obligate receptor mediating CD209-dependent suppression of T-cell activation.

### Anti-CD209 antibodies inhibit CD209 function by competitively disrupting ICAM-2 engagement

ICAM-2, a member of the immunoglobulin superfamily involved in leukocyte trafficking(49), has an incompletely defined role in T-cell biology. To assess its relevance in the tumor microenvironment, we examined ICAM-2 expression on T-cell subsets. Flow-cytometric analysis revealed that ICAM-2 is expressed at significantly higher levels on CD8⁺ than on CD4⁺ T cells in both murine tumor models and human lung adenocarcinoma specimens (Fig. S5D–E), suggesting a preferential role in CD8⁺ T-cell regulation.

We next investigated the mechanism by which CD209-neutralizing antibodies disrupt CD209–ICAM-2 signaling. Structural modeling of the antibody–CD209 interface identified 15 paratope-contact residues, four of which overlap with the ICAM-2–binding footprint (Fig. S5F–G). Alanine scanning of these residues abolished antibody binding, confirming direct engagement of this interface (Fig. S5G–H). Notably, the critical ICAM-2–interacting residue Gly303 is not occupied by the antibody, explaining the partial disruption of CD209–ICAM-2 binding (Fig. 4F, Fig. S5G). Together, these findings indicate that ICAM-2 is preferentially expressed on CD8⁺ T cells and that anti-CD209 antibodies inhibit CD209 function by competitively interfering with the CD209–ICAM-2 interaction, supporting ICAM-2 as the non-redundant receptor mediating CD209-dependent T-cell suppression.

### CD209–ICAM-2 signaling impairs T cell–APC interactions via ROCK1–ERM–mediated mechano-regulation

ERM (ezrin/radixin/moesin) proteins link the plasma membrane to the actin cytoskeleton by binding the cytoplasmic tail of ICAM-2 via their N-terminal FERM domains and anchoring F-actin through their C-terminal domains, thereby regulating cortical rigidity and leukocyte deformability(51–53). Previous studies have shown that ICAM-2 cross-linking induces ERM activation in T cells. Consistent with this, engagement of Jurkat cells with surface-bound hCD209-Fc induced robust ERM phosphorylation at the plasma membrane, an effect that was largely abolished in ICAM-2–deficient cells (Fig. 5A–C), indicating that CD209 activates ERM signaling in an ICAM-2–dependent manner. Given the role of ERM proteins in regulating cortical mechanics, we next assessed T-cell deformability. High-speed centrifugation followed by 3D reconstruction revealed that CD209 engagement significantly increased cellular stiffness, as reflected by a reduced deformability index (DI). In contrast, ICAM-2–deficient cells exhibited comparable deformability under control and CD209-Fc conditions (Fig. 5D–G). Together, these findings demonstrate that CD209–ICAM-2 ligation enhances T-cell cortical rigidity through ERM activation, providing a mechanistic basis for CD209-mediated suppression of T-cell function.

**Figure 5.**
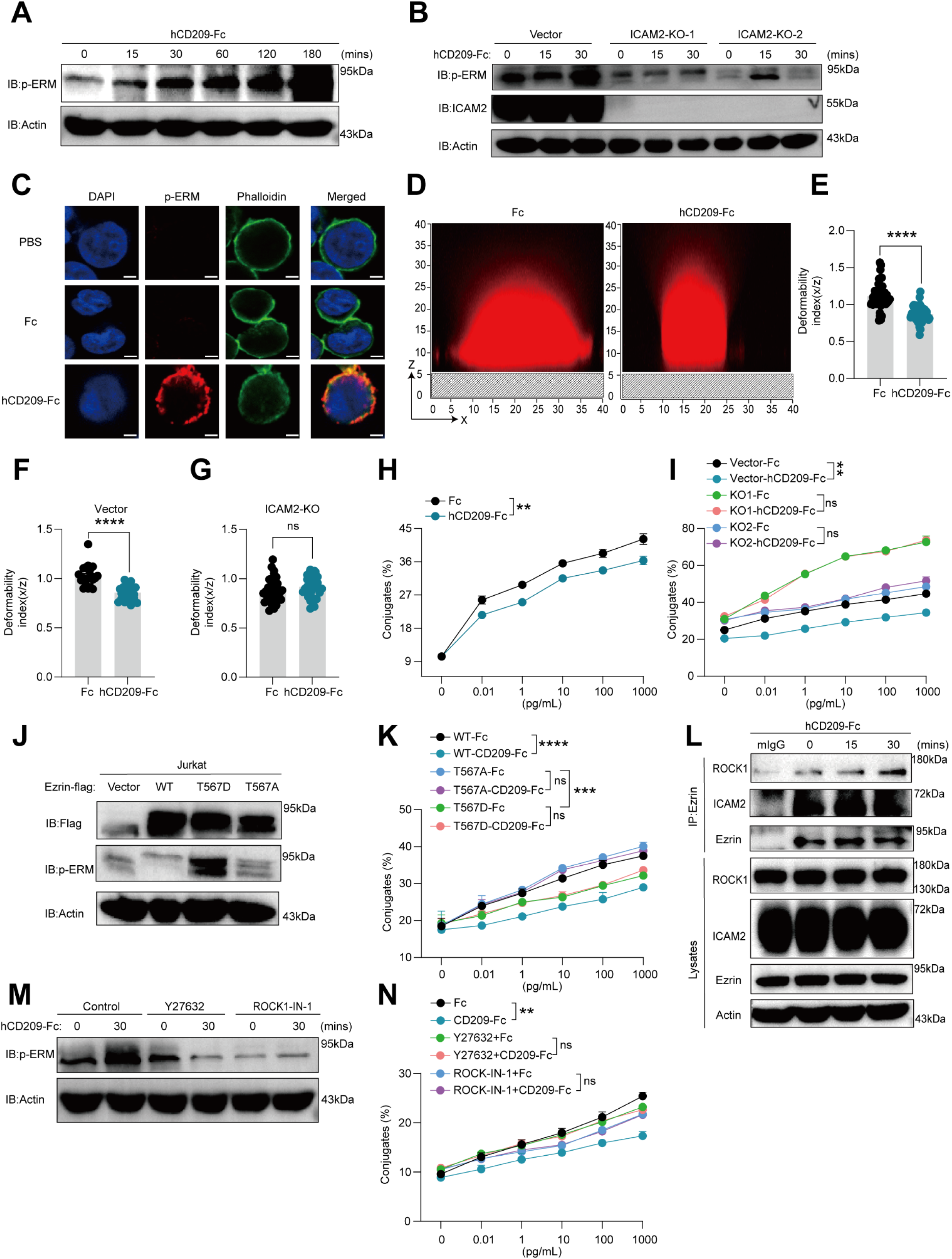
CD209–ICAM-2 signaling impairs T cell–APC interactions via ROCK1–ERM–mediated mechano-regulation. **(A)** Jurkat cells were stimulated with plate-bound Fc or hCD209-Fc for the indicated times, followed by immunoblot analysis of ERM phosphorylation. **(B)** Control or ICAM2-knockout (KO) Jurkat cells were stimulated with hCD209-Fc for the indicated times, followed by immunoblot analysis of ERM phosphorylation. **(C)** Immunofluorescence staining of Jurkat cells treated with Fc or hCD209-Fc for 30 minutes, showing p-ERM (red), F-actin (phalloidin, green), and nuclei (DAPI, blue). **(D–E)** Confocal microscopy analysis of Jurkat cells treated with Fc or hCD209-Fc for 30 minutes. Representative x–z cross-sectional images are shown (D), and the deformability index (x/z) is quantified (E: n=54/56). **(F–G)** Quantification of deformability index in control (F: n=21/21) or ICAM2-KO Jurkat cells (G: n=33/33) following Fc or hCD209-Fc treatment. **(H–I)** Jurkat cells (H), or control and ICAM2-KO Jurkat cells (I), were treated with Fc or hCD209-Fc and co-incubated with SEE-pulsed Raji cells. T cell–APC conjugate formation was quantified by flow cytometry. **(J–K)** Jurkat cells expressing ezrin-WT, phospho-deficient (T567A), or phospho-mimetic (T567D) mutants were analyzed by immunoblotting (J), and their conjugate formation with Raji cells following Fc or hCD209-Fc treatment was quantified (K). **(L)** Immunoprecipitation of ezrin from Jurkat cells stimulated with hCD209-Fc for the indicated times, followed by immunoblot analysis of associated proteins. **(M)** Jurkat cells were pretreated with ROCK inhibitors (Y27632 or ROCK-IN-1) and then stimulated with hCD209-Fc, followed by immunoblot analysis of ERM phosphorylation. **(N)** Jurkat cells pretreated with ROCK inhibitors were stimulated with Fc or hCD209-Fc and co-incubated with SEE-pulsed Raji cells. Conjugate formation was quantified by flow cytometry. Statistical significance was determined using two-tailed unpaired t tests (E–G) and two-way ANOVA (H–I, K, N). *P < 0.05; **P < 0.01; ***P < 0.005; ****P < 0.001. All error bars represent mean ± SEM. Data in ( A-C, H–N) are pooled from two independent experiments. Data in (D-G) are pooled from three independent experiments.

Cortical rigidity is a key determinant of stable T cell–antigen-presenting cell (APC) interactions required for efficient TCR activation(53). We therefore examined whether CD209-mediated increases in T-cell stiffness affect T cell–APC conjugation. Jurkat cells exposed to plate-bound CD209-Fc exhibited a marked reduction in superantigen-dependent APC coupling, whereas ICAM-2–deficient cells maintained normal APC contact following CD209 engagement (Fig. 5H–I, Fig. S5I), indicating that CD209–ICAM-2 ligation destabilizes T cell–APC interactions. This effect was dependent on ERM activation. CD209 significantly impaired APC contact in cells expressing wild-type ezrin, but not in cells expressing either the phospho-deficient ezrin-T567A or the constitutively active ezrin-T567D mutant (Fig. 5J–K), demonstrating a requirement for regulated ezrin phosphorylation.

ROCK1 is a known kinase responsible for ezrin Thr567 phosphorylation(54,55). Consistently, CD209 engagement enhanced ezrin–ROCK1 association (Fig. 5L), and pharmacologic inhibition of ROCK1 abrogated CD209-induced ezrin phosphorylation and restored T cell–APC interactions (Fig. 5M–N). Collectively, these findings define an ICAM-2–ROCK1–ERM signaling axis that establishes a mechanosuppressive checkpoint directly restricting T-cell activation through physical modulation.

### CD209 blockade mediates antitumor immunity independently of Fc effector functions

To determine whether the therapeutic effects of CD209 blockade depend on Fc-mediated cytotoxicity, we generated a humanized anti-CD209 IgG4 antibody (αhCD209-hIgG4; KD = 2.7 × 10⁻⁹ M) lacking Fc effector activity (Fig. S6A–B). Notably, despite elimination of ADCC, αhCD209-hIgG4 fully recapitulated the antitumor efficacy of the parental IgG2a antibody across multiple tumor models. In MC38 tumors, treatment significantly suppressed tumor growth and reduced tumor weight while increasing intratumoral IFN-γ⁺ CD8⁺ T cells (Fig. S6C–E). Similar effects were observed in LLC tumors, with reduced tumor burden and increased IFN-γ⁺ CD4⁺ and CD8⁺ T-cell populations (Fig. S6F–H). Importantly, in immunologically “cold” models, including ID8 ovarian carcinoma and orthotopic GL261 glioblastoma, αhCD209-hIgG4 remained highly effective, reducing ascites accumulation and prolonging survival while maintaining body weight (Fig. S6I–O). Collectively, these findings demonstrate that the antitumor activity of CD209 blockade is independent of Fc-mediated effector functions and is driven primarily by direct inhibition of CD209 signaling.

### CD209 blockade elicits antitumor immunity in human tumor systems and overcomes resistance to PD-1–based immunotherapy

We next assessed the translational potential of CD209 blockade in human tumor systems. Ex vivo treatment of single-cell suspensions from freshly resected lung adenocarcinomas with αhCD209-hIgG4 significantly increased IFN-γ and granzyme B secretion compared with isotype control, indicating reactivation of intratumoral cytotoxic lymphocytes (Fig. 6A–C). In immune-humanized mouse models reconstituted with human PBMCs(9), αhCD209-hIgG4 markedly suppressed tumor growth in both ovarian (OVCAR3) and lung adenocarcinoma (A549) xenografts, accompanied by expansion of intratumoral IFN-γ⁺ CD8⁺ T cells (Fig. 6F–H, Fig. S6P). Notably, in humanized C-NKG models bearing HCT116 (MSS colorectal) or A375 (melanoma) tumors, CD209 blockade demonstrated superior efficacy or strong synergy with PD-1 blockade. It overcame primary resistance to PD-1 blockade in HCT116 tumors (Fig. 6I–K) while enhancing intratumoral CD8⁺ T-cell activation (Fig. 6L), and synergized with PD-1 blockade to further promote tumor regression in A375 tumors (Fig. 6M–O). Residual human myeloid cells persisted within the tumor microenvironment (Fig. S6Q), supporting sustained responsiveness to CD209-targeted therapy. Collectively, these findings demonstrate that CD209 blockade reactivates antitumor immunity in human tumor contexts and overcomes resistance to PD-1–based immunotherapy and further enhances therapeutic responses, supporting its potential as a clinically actionable immunotherapeutic target.

**Figure 6.**
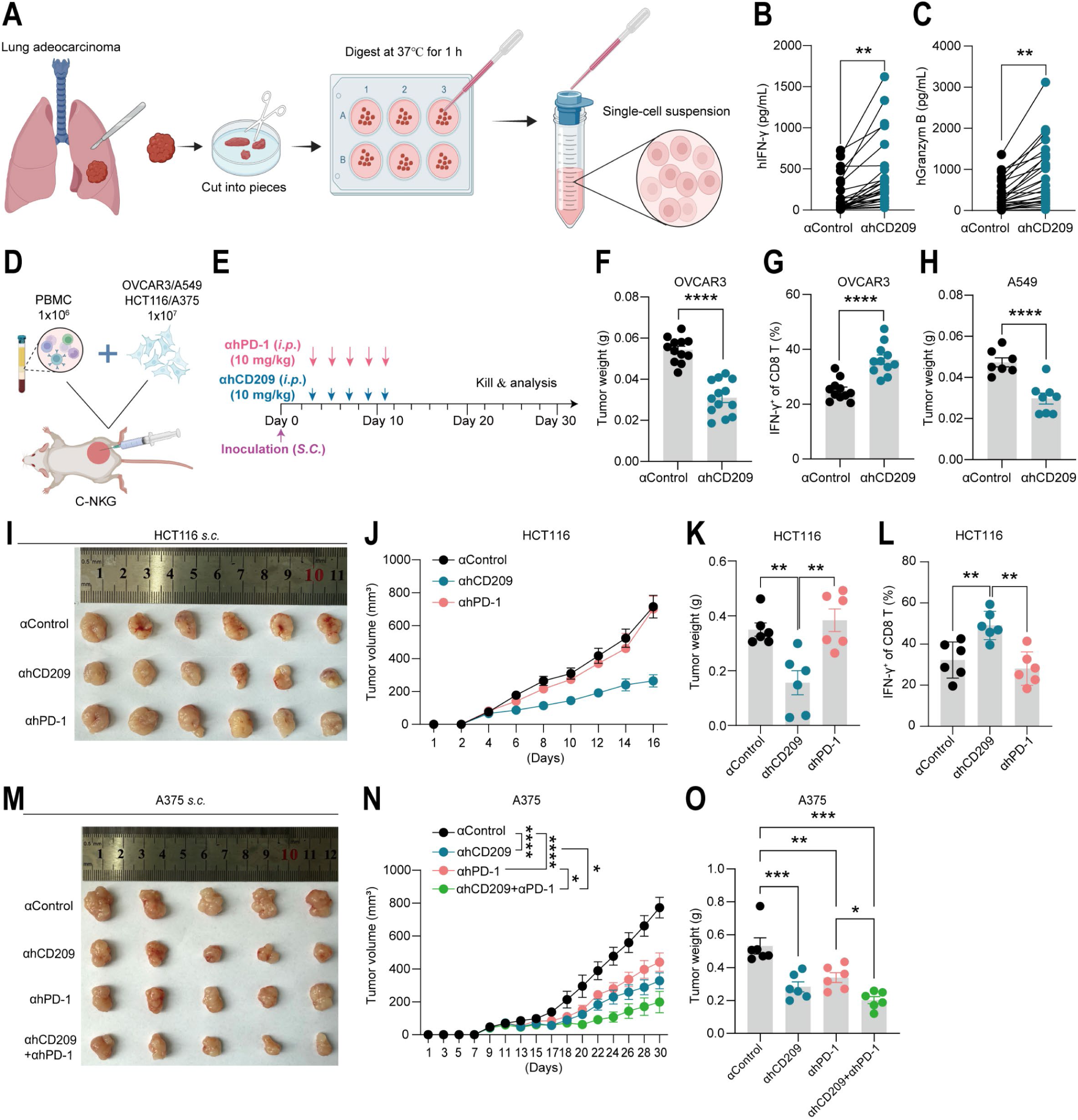
CD209 blockade elicits antitumor immunity in human tumor systems and overcomes resistance to PD-1–based immunotherapy. **(A–C)** Tumor samples from patients with lung adenocarcinoma (LUAD) were dissociated into single-cell suspensions and treated with control hIgG4 or anti-hCD209 hIgG4 for 48 hours (A). IFN-γ (B) and granzyme B (C) levels in the culture supernatant were measured by ELISA. **(D–E)** Schematic of humanized immune system tumor models established by co-engrafting human PBMCs and tumor cells into immunodeficient C-NKG mice. **(F–H)** Antitumor effects of CD209 blockade in humanized models. Tumor weights of OVCAR3 tumors (F: n=12/13), frequency of IFN-γ⁺ CD8⁺ T cells (G), and tumor weights of A549 tumors (H: n=7/8) following treatment with control or anti-hCD209 hIgG4. **(I–L)** Humanized C-NKG mice bearing HCT116 tumors were treated with isotype control, anti-hCD209 hIgG4, or anti–PD-1 (10 mg kg⁻¹; days 2, 4, 6, 8, and 10; n=6 per group). Representative tumor images (I), tumor growth curves (J), tumor weights (K), and the frequency of IFN-γ⁺ CD8⁺ T cells in tumors (L) are shown. **(M–O)** Humanized C-NKG mice bearing A375 tumors were treated with control, anti-hCD209 hIgG4, anti–PD-1, or combination therapy (10 mg kg⁻¹; days 2, 4, 6, 8, and 10; n=6 per group). Representative tumor images (M), tumor growth curves (N), and tumor weights (O) are shown. Statistical significance was determined using two-tailed unpaired t tests (B–C, F–H, K–L, O) and two- way ANOVA (J, N).*P < 0.05; **P < 0.01; ***P < 0.005; ****P < 0.001. All error bars represent mean ± SEM. Data are representative of two independent experiments (A–H) or one independent experiment (I–O).

### CD209⁺ myeloid cells engage T cells in situ and are associated with impaired CD8⁺ T-cell infiltration and poor clinical outcomes

Multiplex immunofluorescence analysis of primary lung adenocarcinoma (LUAD) specimens revealed that CD209⁺ myeloid cells form direct contacts with both CD4⁺ and CD8⁺ T cells (Fig. 7A), with ICAM-2 on CD8⁺ T cells co-localizing at these interfaces, supporting CD209–ICAM-2 interactions in situ (Fig. 7B). Flow cytometric analysis further demonstrated that CD209 is predominantly expressed by tumor-associated macrophages (TAMs), with minimal expression on monocytes and dendritic cells (Fig. S7A–B). Across multiple tumor types, including LUAD, glioblastoma, colorectal carcinoma, and high-grade serous ovarian cancer, quantitative tissue analysis revealed a strong inverse correlation between CD209⁺ TAM density and intratumoral CD8⁺ T-cell abundance (Fig. 7C–F; Fig. S7C–F). Consistent with these observations, pan-cancer survival analyses showed that elevated CD209 expression is significantly associated with worse overall survival across diverse malignancies (Fig. S7G–N). Collectively, these findings identify CD209⁺ myeloid cells as a spatially and clinically relevant immunosuppressive checkpoint that limits CD8⁺ T-cell infiltration and function, thereby promoting tumor immune evasion and adverse patient outcomes.

**Figure 7.**
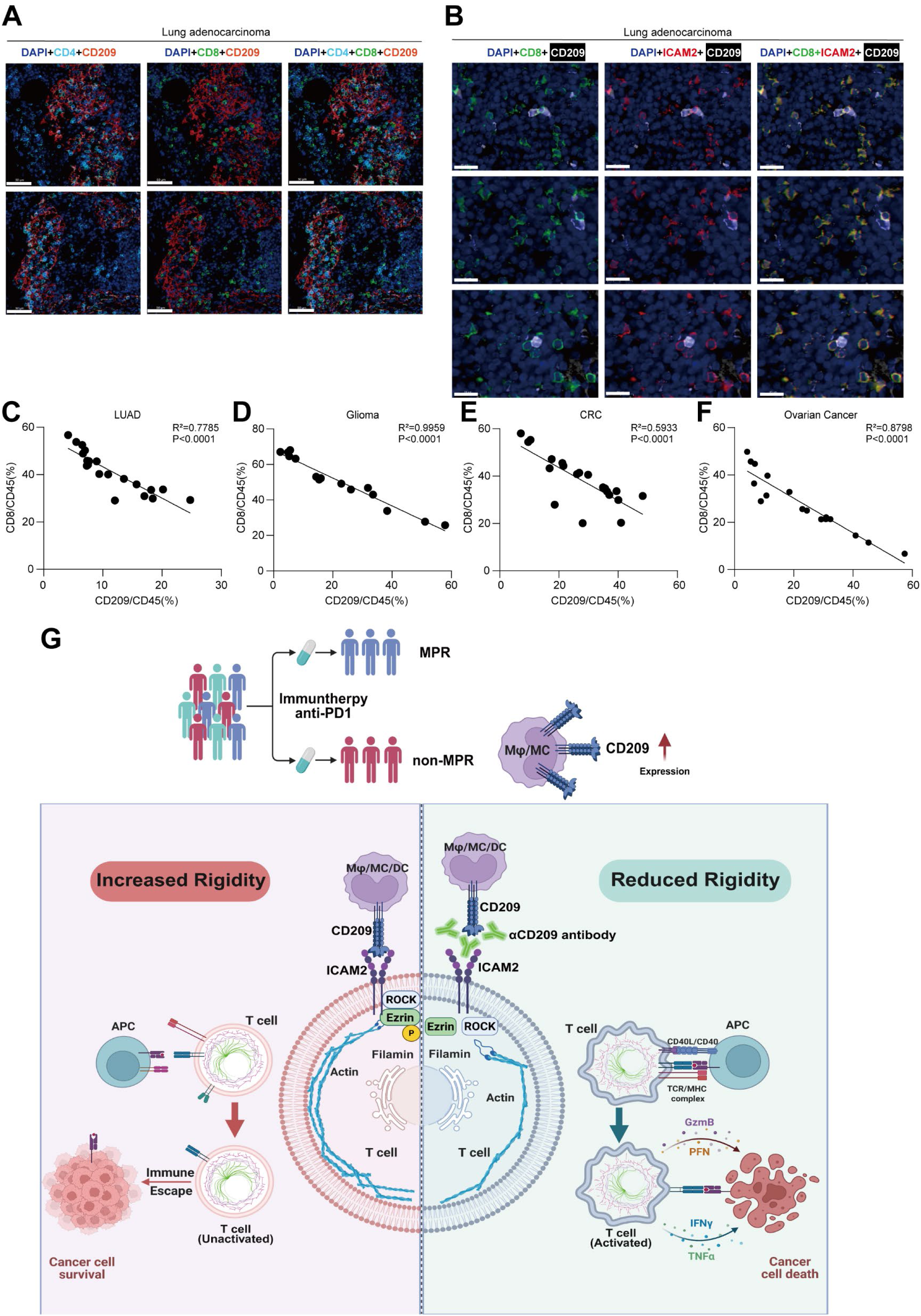
CD209⁺ myeloid cells engage T cells in situ and are associated with impaired CD8⁺ T-cell infiltration. **(A)** Representative multiplex immunofluorescence staining of lung adenocarcinoma tissues showing the spatial distribution of CD209⁺ cells and T-cell subsets. Red: CD209; Green: CD8; Cyan: CD4; Blue: DAPI. Scale bars, 50 μm. **(B)** Representative multiplex immunofluorescence staining of lung adenocarcinoma tissues showing the spatial relationship among CD209⁺ cells, ICAM2⁺ cells, and CD8⁺ T cells. White: CD209; Red: ICAM2; Green: CD8; Yellow: CD8 and ICAM2 colocalization; Blue: DAPI. Scale bars, 20 μm. **(C–F)** Quantification of the relationship between CD209⁺ cells and CD8⁺ T-cell infiltration across tumor types. Correlation analysis between the proportion of CD209⁺ cells and CD8⁺ T cells among CD45⁺ immune cells in lung adenocarcinoma (LUAD, C, n = 20), glioma (D, n = 15), colorectal cancer (CRC, E, n = 20), and ovarian cancer (F, n = 17). Correlations were assessed using Spearman’s rank correlation coefficient. Each dot represents one patient sample. The correlation coefficient (r) and two-sided P value are indicated. **(G)** A sketch of CD209-mediated antitumor immune evasion is shown.

## Discussion

This study identifies a myeloid-derived mechanosuppressive checkpoint that restrains antitumor T-cell immunity through physical modulation of T-cell cortical mechanics. We show that CD209 engages ICAM-2 on T cells to activate a ROCK1–ERM–dependent mechanotransduction pathway, leading to increased cortical rigidity, impaired T cell–antigen-presenting cell interactions, and attenuated TCR signaling. Functionally, disruption of CD209 signaling restores T-cell infiltration and effector function and elicits antitumor immunity across multiple tumor models, including immunologically “cold” and PD-1–refractory tumors. These findings establish CD209 as a mechanistic link between suppressive myeloid cells, T-cell exclusion and resistance to PD-1–based immunotherapy, a major barrier to effective cancer treatment (Fig. 7G)(5,8,56). Consistent with this model, CD209⁺ myeloid-cell infiltration has been associated with poor immunotherapy response and adverse clinical outcomes(29,34). Collectively, our work defines CD209 as a myeloid-driven mechanical immune checkpoint and provides a rationale for targeting the CD209–ICAM-2–ERM axis to overcome T-cell dysfunction and immunotherapy resistance in cancer.

ICAM-2, a member of the immunoglobulin superfamily, is primarily expressed on endothelial cells and platelets and has been implicated in leukocyte adhesion and migration, including LFA-1–dependent transendothelial trafficking(57,58). It has also been reported to promote T-cell survival via AKT signaling(51). In contrast to these established roles, our study identifies ICAM-2 as a previously unrecognized negative regulator of T-cell activation, functioning through a mechanical mechanism that increases cortical stiffness. Although ICAM-2 has not been widely explored as an immunotherapeutic target, prior studies demonstrated that an anti–ICAM-2 monoclonal antibody (EOL4G8) exerts antitumor effects in murine colon carcinoma models in a CD8⁺ T cell–dependent and LFA-1–independent manner(59,60). Moreover, combination therapy with EOL4G8 and IL-12 further enhances tumor regression and is associated with increased ICAM-2 expression on activated CD8⁺ T cells(60). These observations are consistent with our findings and raise the possibility that therapeutic targeting of ICAM-2 may disrupt CD209–ICAM-2–mediated mechanosuppression, thereby restoring effective T cell–APC interactions. This mechanistic link warrants further investigation. Notably, we observed that human CD209 can engage both human and murine ICAM-2, and that neutralizing anti-CD209 antibodies effectively disrupt these interactions across species. These findings support the conservation of the CD209–ICAM-2 axis and validate the use of the hCD209 knock-in mouse model as a physiologically relevant platform for studying CD209-mediated immunoregulation in vivo.

Cellular mechanics has emerged as an important regulator of antitumor immunity, although prior studies have primarily focused on tumor-intrinsic stiffness or extracellular matrix–driven mechanical cues. For example, soft cancer cells enriched in membrane cholesterol can resist T-cell–mediated killing, whereas extracellular matrix stiffening activates ROCK–myosin II–F-actin signaling in tumor cells to suppress dendritic cell maturation and CD8⁺ T-cell priming(61–63). Our study extends this paradigm by showing that mechanical immune regulation can also be imposed directly by myeloid cells onto T cells. Specifically, CD209 expressed by tumor-associated myeloid cells engages ICAM-2 on T cells to activate ERM-dependent cortical remodeling, thereby increasing T-cell stiffness, destabilizing T cell–APC interactions and suppressing T-cell activation. These findings reveal myeloid-driven mechanosuppression as a previously underappreciated mode of immune checkpoint function, distinct from but potentially cooperating with canonical biochemical inhibitory pathways.

Together, these findings reveal a previously unrecognized mode of immune regulation in which myeloid cells impose mechanical constraints on T cells. By linking CD209–ICAM-2 engagement to ERM-dependent cortical stiffening, our study establishes cellular mechanics as an integral component of immune checkpoint function. Targeting this axis may enhance antitumor immunity in PD-1–refractory or T cell–excluded tumors, and T-cell mechanical properties may provide a functional biomarker for patient stratification.

## Materials and Methods

### Ethics statement

Human samples were obtained from Sichuan Provincial People’s Hospital. This study was conducted in accordance with the Declaration of Helsinki and was approved by the Ethics Committee of Sichuan Provincial People’s Hospital, University of Electronic Science and Technology of China. Written informed consent was obtained from all study participants.

All animal experiments were approved by the Institutional Animal Care and Use Committee of Sichuan Provincial People’s Hospital, University of Electronic Science and Technology of China, and were conducted in accordance with institutional guidelines and national regulations for the care and use of laboratory animals.

### Human cancer sample study

A total of 4 LUAD samples, including 2 precancerous normal tissues and matching primary tumor tissues, were included in the scRNA-seq cohort. For histochemical verification, samples of LUAD (n=20) and glioma (n=15), samples of ovarian cancer (n=17), and samples of colon cancer (n=20) were used. All patients were pathologically diagnosed with LUAD, glioma, ovarian cancer or colon cancer and provided informed consent.

We prospectively collected tumor tissue specimens from 41 NSCLC patients receiving combined anti–PD-1 immunotherapy with platinum-based chemotherapy. The cohort comprised 14 responders (favourable clinical response) and 27 nonresponders (suboptimal therapeutic response) to PD-1 blockade therapy. Response categorization was established through the following criteria:

1. **Postoperative immunotherapy cohort:**
2. Patients who received adjuvant immunotherapy demonstrated the following findings: Favourable response:

- Stage II NSCLC patients maintain disease-free survival (DFS) >24 months postimmunotherapy
- Stage IIIa+ patients maintain a DFS >12 months postimmunotherapy Suboptimal Response:
- Disease progression or tumor recurrence within 12 months of immunotherapy initiation
3. Neoadjuvant immunotherapy cohort:
4. Patients receiving preoperative immunotherapy were assessed via the RECIST v1.1 criteria: Favourable response:

- Achievement of Complete Response (CR) or Partial Response (PR)
- Stage II patients maintain a DFS >24 months postimmunotherapy
- Stage III patients maintain a DFS >12 months postimmunotherapy
5. Suboptimal Response:

- Stable disease (SD) or progressive disease (PD) per the RECIST criteria
- Early disease progression/recurrence (≤12 months) despite initial CR/PR

PBMCs were isolated from healthy volunteers. The above human samples were all obtained at Sichuan Provincial People’s Hospital. This study followed the guidelines set forth by the Declaration of Helsinki, and the protocol was approved by the Ethics Committee of Sichuan Provincial People’s Hospital, University of Electronic Science and Technology of China. All study participants signed a written informed consent form.

### Mice

hCD209-KI mice: hCD209-KI mice were purchased from Cyagen Biotech. The gRNA for the mouse *Cd209* gene, the donor vector containing the “human CD209 CDS-3xFLAG tag-3’UTR of the mouse Cd209-rBG pA” cassette, and Cas9 mRNA were coinjected into fertilized mouse eggs to generate targeted knock-in offspring. F0 founder animals, which were bred to wild-type mice to test germline transmission and F1 animal generation, were identified via PCR followed by sequence analysis. F1 generation heterozygous mice were mated to produce homozygous hCD209 knock-in mice. C-NKG mice: C-NKG mice are severely immunodeficient and were independently generated by Cyagen Biotech by knocking out the *Il2rg* gene in the NOD-Scid strain. The C-NKG mouse lacks mature T, B and NK immune cells, with reduced complement activity and a weakened phagocytic effect of macrophages on human cells. It can be used for the efficient transplantation of human hematopoietic stem cells (HSCs), peripheral blood mononuclear cells (PBMCs), or adult stem cells and tissues. All the animal experiments were performed in strict accordance with the relevant ethical guidelines and were approved by the Ethics Committee of Sichuan Provincial People’s Hospital, University of Electronic Science and Technology of China. The mice used in our experiments were housed under specific pathogen-free (SPF) conditions, with an ambient temperature between 20 and 25 °C, humidity between 40 and 70%, and an environmental light/dark cycle of 12 h light/12 h dark.

### Cells

All the cell lines used were from the ATCC. The cell line authentication data obtained through short tandem repeat profiling are publicly available from the ATCC. The cell lines used in this study exhibited different morphologies and growth rates and were routinely monitored for contamination. B16, A549, GL261-LUC, ID8, 293T, EL4, HCT116 and A375 cells were cultured in DMEM supplemented with 10% FBS and 1% penicillin‒streptomycin. 293F cells were cultured in 293F culture medium from Gibco. MC38, LLC, OVCAR3, Jurkat, Raji, and mouse CD4 T cells; mouse CD8 T cells; and primary human single-cell suspensions of tumor tissue were cultured in RPMI-1640 supplemented with 10% FBS, 1% penicillin‒streptomycin and 1% sodium pyruvate.

### Reagents

Phospho-Ezrin (Thr567)/Radixin (Thr564)/Moesin (Thr558) rabbit mAb (clone 48G2, (Cell Signaling Technology Cat# 3726, RRID:AB_10560513)) and the CD102/ICAM-2 rabbit mAb (clone D7P2Q, (Cell Signaling Technology Cat# 13355, RRID:AB_2798188)) were purchased from Cell Signaling Technology. Immunohistochemical antibodies against CD45 (clone PD7/26+2B11, (Maxim biotechnologies Cat# Kit-0024, RRID:AB_3096185)) and CD8 (clone MX117, (Maxim biotechnologies Cat# MAB-1031, RRID: AB_3713025)) were purchased from Maxim. ICAM3 rabbit pAb was purchased from Abconal (cat no. A3923, (ABclonal Cat# A3923, RRID:AB_2765389)). Anti-DC-SIGN (clone 1781, (Abcam Cat# ab218419, RRID:AB_3713027)) was purchased from Abcam. ROCK1 rabbit polyAb (cat no. 21850-1-AP, (Proteintech Cat# 21850-1-AP, RRID:AB_10953526)) was purchased from Proteintech. Anti-HA (clone HA-7, (Sigma-Aldrich Cat# H9658, RRID:AB_260092)) and anti-Flag (clone M2, (Sigma-Aldrich Cat# F1804, RRID:AB_262044)) antibodies for immunoprecipitation were purchased from Sigma. Anti-His-HRP (clone AMC0484, (ABclonal Cat# AE028, RRID:AB_2769867)) was purchased from Abconal. HRP-conjugated goat anti-human IgG (H+L) was purchased from Proteintech (cat no. SA00001-17, (Proteintech Cat# SA00001-17, RRID:AB_2890979)). Phospho-IkBα (Ser32) (clone 14D4, (Cell Signaling Technology Cat# 2859, RRID:AB_561111)) rabbit mAb and IkBα (clone 44D4, (Cell Signaling Technology Cat# 4812, RRID:AB_10694416)) rabbit mAb were purchased from CST. Antibodies against p-ERK (clone 12D4, (Santa Cruz Biotechnology Cat# sc-81492, RRID:AB_1125801)), HSP90 (clone AC-16, (Santa Cruz Biotechnology Cat# sc-101494, RRID:AB_1124018)), actin (clone C-2, (Santa Cruz Biotechnology Cat# sc-8432, RRID:AB_626630)), GAPDH (clone A-3, (Santa Cruz Biotechnology Cat# sc-137179, RRID:AB_2232048)), ERK (G-8, (Santa Cruz Biotechnology Cat# sc-271269, RRID:AB_10611091)), and ezrin (6A84, (Santa Cruz Biotechnology Cat# sc-71082, RRID:AB_1122712)) were purchased from SANTA CRUZ BIOTECHNOLOGY. Antibodies against LCK (ABclonal Cat# A2177, RRID:AB_2764195) and the Flag-tag (ABclonal Cat# AE024, RRID:AB_2769864) were purchased from ABclonal. The anti-p-Lck antibody (Y394) was purchased from R&D Systems (R&D Systems Cat# MAB7500, RRID:AB_2923236). The αhCD209 mAb is secreted by hybridoma cells produced by AtaGenix. The in vivo mAb mouse IgG2a isotype control (clone C1.18.4) was purchased from Selleck (Selleck Cat# A2117, RRID:AB_3662739). The ROCK inhibitors Y-27632 and ROCK1-IN-1 were purchased from MCE (cat nos. HY10071 and HY-Q22471). CellTracker Red CMTPX and CellTracker Blue CMAC were purchased from MCE (cat nos. HY-D1727 and HY-D1462). SEE was purchased from Toxin Technology (cat no: ET404). Anti-mouse CXCL9 (clone MIG-2F5.5) was purchased from Bioxcell (Bio X Cell Cat# BE0309, RRID:AB_2736989). The human IgG4 isotype control (clone R1, (Selleck Cat# A2052, RRID: AB_3713065)), anti-mouse PD-1 (clone RMP1-14, (Selleck Cat# A2122, RRID:AB_3644244)), anti-mouse CD8a (clone 2.43, (Selleck Cat# A2102, RRID:AB_3099521)), and anti-mouse CSF1R (clone AFS98, (Bio X Cell Cat# BE0213, RRID:AB_2687699)). TMZ was purchased from Solarbio (cat no. IT1330). Fluorochrome-conjugated antibodies against mouse CD3 (17A2 (BioLegend Cat# 100221, RRID:AB_2057374)), mouse CD4 (GK1.5, (BioLegend Cat# 100405, RRID:AB_312690)), human CD4 (SK3, (BioLegend Cat# 980802, RRID:AB_2616621)), mouse CD8 (53--6.7, (BioLegend Cat# 100705, RRID:AB_312744)), human CD8 (SK-1, (BioLegend Cat# 980908, RRID:AB_2888883)), mouse CD45 (30--F11, (BioLegend Cat# 103132, RRID:AB_893340)), human CD45 (HI30, (BioLegend Cat# 304014, RRID:AB_314402)), mouse CD11b (M1/70, (BioLegend Cat# 101212, RRID:AB_312795)), mouse F4/80 (BM8, (BioLegend Cat# 123110, RRID:AB_893486)), human CD11b (ICRF44, (BioLegend Cat# 982602, RRID:AB_2922653)), human CD68 (Y1/82A, (BioLegend Cat# 333816, RRID:AB_2562936)), mouse Ly-6C (HK1.4, (BioLegend Cat# 128036, RRID:AB_2562353)), mouse Ly-6G (1A8, (BioLegend Cat# 127628, RRID:AB_2562567)), human CD14 (M5E2, (BioLegend Cat# 982516, RRID:AB_3083241)), human HLA-DR (QA19A44, (BioLegend Cat# 365603, RRID:AB_3083355)), human CD102 (ICAM2-CBR-IC2/2, (BioLegend Cat# 328505, RRID:AB_2295682)), mouse Gr1 (RB6-8C5, (BioLegend Cat# 108424, RRID:AB_2137485)), mouse IFN-γ (XMG1.2, (BioLegend Cat# 505808, RRID:AB_315402)), and human IFN-γ (W19227A, (BioLegend Cat# 383303, RRID:AB_2924583)). A Zombie Violet Fixable Viability Kit was purchased from Biolegend (cat no. 423114). An ELISA MAX^TM^Deluxe Set human IFN-γ was purchased from Biolegend (cat no. 430104). ELISA MAX™ Deluxe Set Mouse IL-2 and ELISA MAX™ Deluxe Set Human IL-2 were purchased from Biolegend (cat nos. 431004 and 431804). A human granzyme B ELISA kit was purchased from CUSABIO (cat no. CSB-E08718h). Ultra-LEAF^TM^-purified anti-mouse CD3ε (clone:145-2C11, (BioLegend Cat# 100359, RRID:AB_2616673)), Ultra-LEAF^TM^-purified anti-mouse CD28 (clone: 37.51, (BioLegend Cat# 102122, RRID:AB_2810331)), Ultra-LEAF^TM^-purified anti-human CD3 (clone: OKT3, (BioLegend Cat# 317348, RRID:AB_2571995)), and Ultra-LEAF^TM^-purified anti-human CD28 (clone: CD28.2, (BioLegend Cat# 302944, RRID:AB_2616668)) antibodies.

### Cell isolation and stimulation

Mouse primary CD4^+^ and CD8^+^ T cells were sorted via the EasySep™ Mouse CD4^+^ T-cell isolation kit (19852, STEMCELL) and the EasySep™ Mouse CD8+ T-cell isolation kit (19853, STEMCELL), respectively. Venous blood was collected into a blood collection tube containing an anticoagulant (heparin sodium). Anticoagulant whole blood was gently mixed with an equal volume of precooled PBS (containing 2% FBS) (diluted 1:1). Ficoll-Paque (half of the diluted blood volume) was placed at the bottom of a 15 mL conical tube. The diluted blood was slowly added along the tube wall to the upper layer of the Ficoll solution (to avoid layer mixing). After density gradient centrifugation, the white film layer (PBMC) was transferred to a new centrifuge tube. After the red blood cells were lysed, they were resuscitated and counted for future use. The sorted T cells, PBMCs or Jurkat cells were stimulated with 2 μg/mL anti-CD3 and anti-CD28 antibodies in 96-well plates (1 × 10^5^ cells per well) for ELISA and stimulated with 2 μg/mL anti-CD3 and anti-CD28 antibodies in 12-well plates (1 × 10^6^ cells per well) for immunoblot analysis and flow cytometric analysis.

### Recombinant protein expression and purification

Most membrane proteins in eukaryotes usually undergo glycosylation to perform their biological functions. Therefore, we purified the CD209 and ICAM2 proteins via a eukaryotic protein expression system to construct Fc fusion expression vectors or His tag expression vectors (the skeletal vector was pINFUSE_hIgG2_Fc2). 293F cells were cultured, and the optimal growth density of these cells was 1 × 10^6^ cells/mL. The density of the 293F cells was maintained at 1 × 10^6^ cells/mL the day before transfection. On the second day, when the cell density reached 2 × 10^6^ cells/mL, the target plasmid was transfected using Polyplus transfection reagent. The cell viability was observed 7–8 days after transfection. When the viability was approximately 30%, the cell culture supernatant was collected. The protein with the Fc tag was purified via Protein G, and the protein with the His tag was purified via a nickel column. Following clarification by centrifugation, the supernatant was subjected to affinity chromatography using protein G resin (for Fc-tagged proteins) and Ni-NTA resin (for His-tagged proteins). The immobilized hCD209/hICAM2-Fc fusion protein was eluted via stepwise pH gradient elution using 0.1 M glycine-HCl buffer (pH 3.0) followed by immediate neutralization with 1 M Tris-HCl (pH 8.5) at a 1:9 (v/v) ratio. Concurrently, the hCD209/hICAM2-His recombinant protein underwent immobilized metal affinity chromatography (IMAC) purification with sequential washes: nonspecific binding removal using 10 mM imidazole in equilibration buffer (20 mM phosphate, 500 mM NaCl, pH 7.4), followed by isocratic elution with 250 mM imidazole in elution buffer. Both purified protein fractions were buffer-exchanged into PBS (pH 7.4) via PD-10/30 desalting columns, concentrated to 1 mg/mL via Amicon® Ultra centrifugal filters (10/30 kDa MWCO), and cryopreserved at −80 °C.

### Flow cytometry analysis of protein‒cell binding

Primary mouse CD4^+^ and CD8^+^ T cells were stimulated with 2 μg/mL anti-CD3 and anti-CD28 antibodies for 24 hours at 37 °C. Jurkat cells were stimulated with 2 μg/mL anti-CD3 and anti-CD28 antibodies for 12 hours at 37 °C. The activated cells were subsequently equally divided into two groups. For each group, 2 μg of Fc or hCD209-Fc was introduced into the cells. The binding reaction was carried out in FACS buffer containing 2 mM Ca^2+^ without EDTA. The mixture was incubated for 30 minutes on a shaker at 4 °C. After washing to remove unbound components, fluorescein isothiocyanate (FITC)-labelled fluorescent secondary antibodies were applied for staining at 4 °C for 30 minutes. Ultimately, flow cytometry was employed to detect binding events.

### Anti-hCD209 monoclonal antibody production

The anti-hCD209 monoclonal antibodies were produced and purified through the following procedures (1) Immunization: 50 μg of recombinant hCD209-ECD-Fc protein was emulsified in 100 μL of phosphate-buffered saline (1 × PBS) with 100 μL of Freund’s complete adjuvant (Sigma F5881). Six-to eight-week-old male C57BL/6 mice (Wuhan AtaGenix) were subcutaneously injected at 21-day intervals for four immunizations. (2) Splenocyte isolation: Mechanically dissociate splenic tissue through 70 μm cell strainers. The erythrocytes were lysed with 10 mL of ACK lysis buffer (Beyotime C3702) for 10 min at 25 °C, followed by neutralization with 40 mL of serum-free DMEM and filtration. (3) Cell fusion: Splenocytes were combined with SP2/0 myeloma cells at a 1:5 ratio and centrifuged at 500 × g for 5 min. The pellets were resuspended in 1 mL of 50% PEG-1500 (Roche), incubated for 90 sec, and then gradually diluted with 50 mL of serum-free medium. After 10 min of incubation, the mixture was centrifuged and resuspended in 10 mL of HAT selection medium. (4) Plating: Seed the cell suspension at 1–3 × 10⁴ cells/well in 96-well plates containing 280 μL of HAT medium. The cells were maintained at 37 °C in a 5% CO₂ humidified incubator for 10 days. (5) Primary screening: ELISA plates were coated with 1 μg/mL hCD209-Fc overnight at 4 °C. The hybridoma supernatants were screened via a standard ELISA protocol. (6) Subcloning: Positive wells whose OD450 > 2.0 × the negative control were expanded. (7) Secondary screening: 100 μL of the supernatant was retested after 48 hours of culture with serial dilutions (1:2 to 1:16) against hCD209-Fc-coated plates. (8) Clone selection: Identify antigen-specific clones demonstrating dose-dependent binding (OD450 inversely proportional to the dilution factor).

### Screening of neutralizing anti-hCD209 mAbs

The neutralizing anti-hCD209 mAb was produced and screened as described previously(13,64). To further screen out functional monoclonal antibodies that block human CD209. 2 μg/mL anti-CD3 and anti-CD28 antibodies were coated overnight at 4 °C in high-affinity 96-well plates, and then, hCD209-Fc (10 μg/mL in 100 μL of 1×PBS) was added to the plates at room temperature for 2–4 hours. After the PBS was aspirated, Jurkat cells, EL4 cells, mouse CD4^+^ T cells, or mouse CD8+ T cells (1 × 10^5^ cells per well) were added. Finally, the control or anti-hCD209 hybridoma culture supernatant was added. The cells were cultured at 37 °C for 48 hours, the culture supernatant was collected, and the level of the cytokine IL-2 was detected via ELISA. The monoclonal hybridoma that could restore the inhibition of IL-2 secretion by hCD209-Fc was the CD209 functional blocking antibody we needed. The monoclonal hybridoma cells were resuscitated, and a large amount of antibody purification was carried out for subsequent functional verification in mice.

### Humanized anti-hCD209 monoclonal antibody production and purification

The humanized anti-hCD209 monoclonal antibodies developed through CDR grafting were produced and purified via the following procedures: (1) Sequence comparison: Sequences were analysed via https://www.ncbi.nlm.nih.gov/igblast and https://www.abysis.org/to select appropriate structural prediction models. (2) Residue analysis: Key framework residues in the VH and VL domains were evaluated on the basis of human germline prevalence to identify amino acids potentially affecting the CDR conformation. (3) Backmutations: Selected framework residues that might influence antibody‒antigen interactions are reverted to their murine counterparts. (4) Vector construction and expression: Humanized antibody variants were cloned and inserted into the pcAGGS-hIgG4 expression vector and transiently expressed in 293T cells. (5) Antibody screening: Culture supernatants were harvested 48 hours posttransfection. The binding affinities between hCD209 and humanized antibodies were quantified by surface plasmon resonance (SPR), with the highest-affinity clone selected for further development. (6) Scale-up production: Large-scale transfection of 293F cells was performed via VH and VL expression plasmids, followed by antibody harvesting and purification for downstream applications.

### Mouse tumor implantation and treatments

All the animal experiments were performed in strict accordance with the relevant ethical guidelines and were approved by the Department of the Ethics Committee of Sichuan Provincial People’s Hospital, University of Electronic Science and Technology of China. For the in vivo tumor growth experiments, a suspension containing 1 × 10⁶ MC38 or 1 × 10⁶ LLC cancer cells were inoculated subcutaneously into the right flank region of hCD209-KI mice. Beginning on day 5 after inoculation, the tumor sizes were recorded every 2 days via a Vernier calliper and calculated with the formula V = L×W²×0.5, where V represents the tumor volume, L denotes the maximum longitudinal diameter, and W signifies the corresponding perpendicular diameter. Beginning on day 7 after inoculation, the mice bearing tumors were randomly divided into different groups and subjected to intraperitoneal administration of αControl mIgG, αhCD209 mIgG, αControl hIgG4, or αhCD209 hIgG4 at a dose of 10 mg/kg per mouse on days 7, 9, 11, 13, and 15 after tumor implantation, respectively. As the tumor mass progressed to a predetermined size, the mice were humanely euthanized following a specific protocol, and the tumor tissues were excised and collected for subsequent experimental analyses.

The ID8 ovarian cancer model was established via the intraperitoneal injection of 3 × 10^6^ ID8 ovarian cancer cells. Beginning on day 7 after inoculation, the mice bearing tumors were randomly divided into different groups and subjected to intraperitoneal administration of αControl mIgG, αhCD209 mIgG, αControl hIgG4, or αhCD209 hIgG4 at a dosage of 10 mg/kg per mouse on days 7, 9, 11, 15, 19, and 21 after tumor implantation, respectively. After 6 consecutive administrations, the occurrence of ascites in the mice was observed. On the 30th day, the mice were sacrificed after dislocation, and serum and ascites were collected for subsequent studies. The mice were subjected to humane euthanasia through a standardized procedure. They were placed in a designated euthanasia chamber, and CO₂ euthanasia was performed by gradually introducing carbon dioxide into the chamber over 10 minutes to ensure a peaceful transition. Following CO₂ euthanasia, each mouse underwent cervical dislocation to confirm irreversible loss of consciousness and death, according to accepted veterinary guidelines.

### Establishment of an orthotopic glioblastoma model

GL261-LUC cells (1 × 10^5^) were implanted into the striatum of the brain at a depth of 2.5 mm to establish an orthotopic glioblastoma-bearing model. The GL261-LUC model mice were randomly divided into four groups as follows: (1) Control group (the mice were intravenously injected with 500 μg of control mAb on days 5, 7, 10 and 15 and sonicated with focus ultrasound after 1 h); (2) TMZ group (the mice were intraperitoneally injected with 100 mg/kg temozolomide on days 6, 8, 10, 12, 14, and 16); (3) αhCD209 mAb group (the mice were intravenously injected with 500 μg/mouse of αhCD209 mAb on days 5, 7, 10 and 15 and sonicated with focus ultrasound after 1 h); and (4) TMZ + αhCD209 mAb group (the mice were intravenously injected with 500 μg of αhCD209 mAb on days 5, 7, 10 and 15 and sonicated with focus ultrasound after 1 h. The mice were also injected with 100 mg/kg temozolomide on days 6, 8, 10, 12, 14, and 16). The tumors established from the GL261-LUC cells were monitored with an IVIS Spectrum system (PerkinElmer) after the injection of D-luciferin potassium salt (75 mg/kg).

### Ultrasound-mediated BBB opening

A US transducer (0.5 MHz and 30 mm diameter) driven by a function generator connected to a power amplifier was used to open the BBB of mice bearing orthotopic glioblastoma-bearing tumors. A detachable water-filled cone served as both an acoustic coupler and a beam guide to direct focused ultrasound into the brain. The acoustic parameters included a peak negative pressure of 0.6 MPa, a pulse repetition interval of 1 ms, a fundamental frequency of 0.5 MHz, and a total sonication duration of 90 s per target. A total of 50 μL of microbubbles was injected intravenously into the mice before sonication.

### Detection of αhCD209 enrichment at the GL261 tumor site

To determine the enrichment of the αhCD209 mAb at the tumor site in the mice bearing GL261 orthotopic GBM, the αhCD209 mAb was modified with FITC. After intravenous injection of 500 μg of αhCD209 mAb-FITC for 1 h, the mice were sonicated with focused ultrasound. The concentration of αhCD209 mAb-FITC enriched at the tumor site was monitored with an IVIS Spectrum (PerkinElmer) after 30 min.

### Single-cell sequencing

Precancerous tissues and primary cancerous tissues from patients with lung adenocarcinoma were collected. These tissues were washed with ice-cold PBS and cut into small pieces, which were then subjected to enzymatic digestion with collagenase D and DNase I via a gentleMACS Tissue Dissociator (Miltenyi). The cells were filtered through a 40 μm filter and washed with PBS. Viability tests (above 95%) and quality control were conducted on the cells. Using the 10x Genomics ChromiumTM system, GelBeads with sequence labels, sample and reagent premix solutions, and oil were loaded into their respective injection channels. Through the “T-shaped channels” formed by the microfluidic channel network, single-cell microreaction system GEMs wrapped by oil droplets finally formed. After anti-transcription by GEMs, barcode-labelled cDNA is formed. The standard sequencing DNA library was established through PCR, enzyme digestion and fragment optimization. High-throughput sequencing was performed on the established libraries via the Illumina platform. The raw data were submitted to the Gene Expression Omnibus database under accession number GSE298300. MC38 subcutaneous tumors were established in hCD209-KI mice. Beginning on day 7 after inoculation, the mice bearing tumors were randomly divided into different groups and subjected to intraperitoneal administration of αControl hIgG4 or αhCD209 hIgG4 at a dose of 10 mg/kg per mouse on days 7, 9, 11, 13, and 15 after tumor implantation. On the 16th day, the mice were subjected to humane euthanasia through a standardized procedure. The samples were washed with Hanks’ balanced salt solution (HBSS) three times, minced into small pieces, and then digested with 3 mL of CelLiveTM Tissue Dissociation Solution (Singleron) with a Singleron PythoN™ Tissue Dissociation System at 37 °C for 15 min. The cell suspension was collected and filtered through a 40-micron sterile strainer. Afterward, the GEXSCOPE® red blood cell lysis buffer (RCLB, Singleron) was added, and the mixture [cell:RCLB=1:2 (volume ratio)] was incubated at room temperature for 5–8 min to remove red blood cells. The mixture was then centrifuged at 300 × g and 4 °C for 5 min to remove the supernatant, after which the mixture was gently suspended in PBS. Finally, the samples were stained with Trypan blue, and the cell viability was evaluated microscopically. Single-cell suspensions (2 × 10^5^ cells/mL) with PBS (HyClone) were loaded onto a microwell chip via the Singleron Matrix^®^Single Cell Processing System. Barcoding beads are subsequently collected from the microwell chip, followed by reverse transcription of the mRNA captured by the barcoding beads to obtain cDNA and PCR amplification. The amplified cDNA was then fragmented and ligated with sequencing adapters. The scRNA-seq libraries were constructed according to the protocol of the GEXSCOPE^®^ single-cell RNA library kit (Singleron). Individual libraries were diluted to 4 nM, pooled, and sequenced on DNBSEQ-T7 with 150 bp paired-end reads. The raw data were submitted to the Gene Expression Omnibus database under accession number GSE308465.

### Flow cytometry analysis of the tumor microenvironment

Human tumors and mouse tumors were collected, dissociated mechanically, and digested with 2 mg/ml collagenase IV (Sigma) and 0.2 mg/ml DNase I (Biofroxx) in serum-free DMEM at 37 °C. After 40 min, the enzyme activity was neutralized by the addition of cold RPMI-1640/10% FBS, the tissues were passed through 70 μM cell strainers (Biologix Group Limited), and single-cell suspensions in T-cell culture medium (RPMI-1640, 10% FBS, 100 IU/ml penicillin, 100 μg/ml streptomycin, 0.5% β-mercaptoethanol) were stimulated with a cell activation cocktail (with brefeldin A) for 3 h (for intracellular staining). After stimulation, the cells were incubated with anti-CD16/CD32 (BioLegend) or Human TruStain FcX™ before being stained with fluorochrome-conjugated monoclonal antibodies. Cell surface staining was performed for 30 min at 4 °C. Intracellular staining was performed via a fixation/permeabilization kit (BioLegend). A Zombie Violet Fixable Viability Kit (1:800; Biolegend) was used to exclude dead cells. Flow cytometry data analysis was performed by using CytExpert. The specific gating strategies used for immune cells via flow cytometry are detailed in supplementary data 3, 4 and 5.

### ELISA

The culture supernatants of Jurkat cells, mCD4^+^ T cells, mCD8^+^ T cells, EL4 cells or PBMCs were used for detection of the cytokine IL-2. The orbital blood of the tumor-bearing mice was collected. After standing still, the serum was collected for subsequent cytokine detection. MC38 or B16 tumors (tumors were cut into small pieces of ∼50 mm^3^ per piece, 3–4 pieces per 48-well plate) were cultured in 0.5 mL of RPMI medium supplemented with 10% fetal bovine serum (FBS) and antibiotics overnight. The tumor supernatant and serum were collected, centrifuged and analysed via IFN-γ or CXCL9 ELISA. The cytokine concentration was normalized to the weight of the tumors in each well.

### In vivo CD8^+^ or macrophage depletion experiments

To deplete CD8+ T cells and macrophages within mouse tumors, αCD8+ and αCSF1R- neutralizing antibodies were used. Following the subcutaneous inoculation of MC38 tumor cells, the αCD8 antibody was administered at 200 μg per mouse every other day beginning on day one and continuing throughout the experiment. For macrophage depletion, an αCSF1R antibody was administered to the mice one week before MC38 tumor inoculation. The mice received 400 μg of αCSF1R antibody every other day, with dosing continuing after tumor inoculation until the experiment was concluded. After the experiment, tumor tissues were harvested from the mice, and flow cytometry was performed to evaluate the efficacy of CD8^+^ T-cell and macrophage depletion within the tumor microenvironment.

### In vivo anti-PD-1 treatment experiments

All the animal experiments were performed in strict accordance with the relevant ethical guidelines, and the protocols were approved by the Animal Care and Use Committee at Sichuan Provincial People’s Hospital, University of Electronic Science and Technology of China. Each individual hCD209-KI mouse (female, 8 weeks old) was randomized into one of four groups (n=12 for each group) and injected with 1 × 10⁶ MC38 cells subcutaneously in the right flank. Starting on day 7 after inoculation, we treated one group of model mice with 10 mg/kg control mouse IgG, one group with 10 mg/kg αhCD209 mAb, one group with 10 mg/kg anti-PD-1 mAb and one group with 10 mg/kg αhCD209 mAb and 10 mg/kg anti-PD-1 mAb from the seventh day after inoculation of the tumor cells every two days, five times in total. In this study, when MC38 tumors had grown for 18 days, all the mice were sacrificed, the tumors were removed and weighed for pretreatment, and further experiments were conducted.

### Mass spectrometry

To determine the functional receptor of CD209 residing on the T-cell surface, we initiated the activation of Jurkat T cells by employing antibodies specific to hCD3 and hCD28 and incubating them at 37 °C for 8–10 hours. Subsequently, 10 dishes (10 cm) of activated Jurkat T cells were harvested, and membrane plasmic separation was performed, after which the membrane proteins were collected and evenly distributed into two distinct groups. The groups were subjected to binding with Fc or hCD209-Fc at a concentration of 30 μg/mL, with the binding process carried out overnight on a shaker maintained at 4 °C. Next, protein G beads were introduced to enrich Fc and hCD209-Fc, and the immunoprecipitation (IP) technique was applied to isolate the membrane proteins bound to CD209.

Then, elution was performed using 0.3% SDS on a shaker at 4 °C for 4 hours. The resulting eluent samples underwent reduction and alkylation processes utilizing dithiothreitol (DTT) and iodoacetamide and were subsequently digested with trypsin overnight. Trypsin is usually used to enzymatically cleave proteins, and this process breaks down complete proteins into smaller peptide segments. The digested peptide segment samples are dissolved in an appropriate buffer solution, such as an aqueous solution containing acetonitrile and formic acid, and then ionized through an electrospray device. After charged protein or peptide ions enter the mass spectrometer, the mass spectrometer separates and detects them on the basis of their mass‒charge ratio. The mass spectrometer records the signal intensity of ions with different mass‒to‒charge ratios and generates a mass spectrum. The mass spectrometry data were compared with the protein database. For peptide mass spectrometry data, the measured peptide mass is matched with the theoretical peptide mass of known proteins in the database through search engines (such as Mascot and SEQUEST). On the basis of the matching results, the proteins present in the sample can be identified.

### *In vitro* protein‒protein interaction experiments

Immunoprecipitation assays were conducted to explore the interaction between hCD209-Fc and hICAM2-his proteins. Initially, 2 μg of the Fc and hCD209-Fc proteins were separately incubated with Protein G agarose beads on a 4 °C shaker for 1 hour. Unbound proteins were removed by washing with IP buffer. Then, 2 μg of hICAM2-his protein was added to each group and incubated on a shaker at 4 °C for 3 hours. The beads were washed four times with IP buffer to eliminate nonspecific binding. After the mixture was decanted, 100 μL of 1 × loading buffer was added, and the samples were boiled at 95 °C for 5 minutes. Western blotting (WB) was performed to detect potential interactions between the two proteins.

To further verify whether the CD209 monoclonal antibody could block the binding of CD209 to its receptor ICAM2 in vitro, 2 μg of hICAM2-his protein was incubated with Ni beads on a shaker at 4 °C for 1 hour. Unbound proteins were washed away with IP buffer. The beads were then evenly divided into four groups. Fc, hCD209-Fc, hCD209-Fc + αControl mAb, and hCD209-Fc + αhCD209 mAb were added to each group. After incubation on a 4 °C shaker for 3 hours, the beads were washed four times with IP buffer. The liquid was removed, 100 μL of 1 × loading buffer was added, and the samples were boiled at 95 °C for 5 minutes. WB was conducted to assess whether the CD209 monoclonal antibody could inhibit the binding of CD209 to ICAM2.

### Lentivirus-mediated gene knockout in Jurkat cells

We transfected the lentiviral construct Lenti-Crispr-ICAM2/ICAM3-gRNA-GFP or the empty vector with VSV-G and PAX2 into HEK293T cells via PEI according to the manufacturer’s protocol. We collected viral supernatants after 48 h and 72 h and infected the Jurkat cells. One week post infection, single GFP-positive cells were isolated and individually allocated into 96-well culture plates via a Beckman flow cytometer. After a subsequent two-week culture period, the cell lines were transferred to 6-well culture plates for further expansion. When the cells grew to a certain density, the effect of gene knockout was verified via Western blotting as described previously(65).

### Immunoblotting and immunoprecipitation

The cells were harvested and lysed on ice in lysis buffer containing 0.5% Triton X-100, 20 mM HEPES (pH 7.4), 150 mM NaCl, 12.5 mM β-glycerophosphate, 1.5 mM MgCl2, 10 mM NaF, 2 mM dithiothreitol, 1 mM sodium orthovanadate, 2 mM EGTA, 20 mM aprotinin, and 1 mM phenylmethylsulfonyl fluoride for 30 minutes, followed by centrifugation at 12,000 rpm for 15 minutes to extract clear lysates(66). Briefly, the cell lysates were incubated with 1 μg of antibody at 4 °C overnight, followed by incubation with A-Sepharose or G-Sepharose beads for 2 hours. The beads were subsequently washed four times with lysis buffer, and the precipitates were eluted with 2× sample buffer. The eluates and whole-cell extracts were resolved by SDS‒PAGE followed by immunoblotting with antibodies.

### Alpha Fold 3 prediction of structural analysis

Structural prediction. Structural models of protein‒protein interactions were predicted via alphafold3 (https://alphafoldserver.com/welcome). Protein interaction interfaces were visualized and analysed via PyMOL software.

### Cellular rigidity measurements

Cellular rigidity was measured according to a previous report(53). Briefly, confocal glass-bottom cell culture dishes were precoated with 100 μg/mL poly-L-lysine at 4 °C overnight. The dishes were subsequently coated with Fc or hCD209-Fc proteins at room temperature for 2 hours. Jurkat T cells were prestained with CellTracker Red CMTPX in serum-free medium for 30 minutes. Then, 2 × 10^5^ stained Jurkat T cells were distributed into each confocal glass-bottom cell culture dish coated with Fc or hCD209-Fc proteins. After incubation at 37 °C for 30 minutes to allow binding and stimulation, the cells were fixed with 2% paraformaldehyde. Finally, all the samples were subjected to centrifugation at 2600 × g for 10 minutes. Coverslips were then washed with PBS, dried and mounted on slides with FluorSave mounting medium (Calbiochem). The samples were visualized with a Zeiss LSM 900 microscope (Leica Microsystems) with a 40 × water mirror objective, and z sections separated by 0.3 µm were acquired. X-Z reconstruction of the image was carried out via the Zeiss LSM 900 microscope confocal microscope software system. A box around the X-Z representation obtained on the equatorial plane of each cell was drawn, and the image was quantified via the software provided by Zeiss LSM 900 microscope confocal microscopes (x corresponds to the width of the cell, and z corresponds to the thickness of the cell). Calculations of the X-Z ratio were performed with a Kaleidagraph and used as the cellular deformability index.

### Analysis of T-cell‒APC conjugate formation

T-cell‒APC conjugate formation was analysed according to a previous report(53). Briefly, high-affinity 24-well plates were coated with Fc and hCD209-Fc proteins at room temperature for 2 hours. Then, 1 × 10^6^ Jurkat T cells stably expressing GFP were seeded in each well and stimulated at 37 °C for 30 minutes. While stimulating the proteins, the Raji cells were loaded with Cell Tracker Red CMTMR (molecular probe) and pulsed with the mixed superantigen SEE at 37 °C for 15 minutes at different concentrations. T cells and APCs were mixed at a 1:1 ratio, centrifuged at 18 × g for 10 seconds, and incubated at 37 °C for 25 minutes. After mild resuscitation, the formation of the conjugate was analysed by flow cytometry via a FACScan flow cytometer (Beckman). The percentage of the conjugate is the GFP^+^CMTMR^+^ cell count/(GFP^+^CMTMR^+^ cell count + GFP^+^CMTMR^-^ cell count) × 100.

### Establishment of immune system humanized mouse cancer models

All the animal experiments were performed in strict accordance with the relevant ethical guidelines and were approved by the Animal Care and Use Committee at Sichuan Provincial People’s Hospital, University of Electronic Science and Technology of China. After one week of adaptation to the environment, each individual C-NKG mouse (female, 6 weeks old) was randomized into two groups and injected with 1 × 10^7^ pretreated OVCAR3, A549, HCT116 or A375 cells with 1.0 × 10^6^ PBMCs subcutaneously in the right flank. Beginning on Day 3 for OVCAR3, A549, HCT116 and A375 cells after tumor implantation, the tumor sizes were recorded every day via a Vernier calliper. The tumor volume was determined via the formula V = L×W²×0.5, where V represents the tumor volume, L denotes the maximum longitudinal diameter, and W signifies the corresponding perpendicular diameter. Beginning on day 3 after inoculation, tumor-bearing c-NKG mice were peritoneally administered 10 mg/kg/human IgG4 isotype control, anti-hCD209 hAbs, anti-hPD-1 hAb, or anti-hCD209 hAbs with anti-hPD-1 hAb every other day, five times in total. On the day after the last administration, the mice were subjected to humane euthanasia through a standardized procedure, and the tumor tissues were resected, weighed and processed for flow cytometric analysis.

### In vitro αhCD209 hIgG4 treatment of a single-cell suspension of human lung adenocarcinoma

Fresh human lung adenocarcinoma samples were collected, dissociated mechanically, and digested with 2 mg/ml collagenase IV (Sigma) and 0.2 mg/ml DNase I (Biofroxx) in serum-free DMEM at 37 °C. After 40 mins, the enzyme activity was neutralized by the addition of cold RPMI-1640/10% FBS, and the tissues were passed through 70 μM cell strainers (Biologix Group Limited) and single-cell suspensions in T-cell culture medium (RPMI-1640, 10% FBS, 100 IU/ml penicillin, 100 μg/ml streptomycin, 0.5% β-mercaptoethanol). The single-cell suspensions were divided into two equal parts. In accordance with the volume of the resuspended medium, αControl hIgG4 or αhCD209 hIgG4 at a final concentration of 30 μg/mL was added. The cells were incubated in a cell culture incubator at 37 °C for 48 hours. The culture supernatant was collected, and the contents of interferon-γ (IFN-γ) and granzyme B in the supernatant were detected via an enzyme-linked immunosorbent assay (ELISA) kit.

### Multicolor fluorescent immunohistochemical staining

The paraffin-embedded tissue sections were dewaxed with xylene until rehydration was achieved. Subsequently, antigen retrieval was conducted via thermal treatment in sodium citrate buffer at 95 °C to adequately expose the antigenic determinants. To eliminate endogenous peroxidase activity that could lead to nonspecific staining, the sections were treated with 3% hydrogen peroxide. Nonspecific protein binding sites were blocked by incubating the sections with 5% bovine serum albumin (BSA) at room temperature for 30 minutes. The specific primary antibody was incubated overnight at 4 °C to allow specific antigen‒antibody binding under optimal conditions. After thorough washing, the corresponding secondary antibody conjugated with a fluorescent dye was added, and the mixture was incubated at room temperature in the dark for 50 minutes to ensure efficient signal amplification. The fluorescent dye reaction was carried out for 10–15 minutes to facilitate the binding of the dye to the antibody, thereby generating a detectable fluorescent signal. Following three washes with phosphate-buffered saline (PBS) to remove unbound dye, the process of antigen retrieval was repeated for the sequential staining of the second and third fluorescent markers, with identical incubation and reaction steps to ensure uniform signal generation across all the markers. After the final staining, the slides were placed in PBS (pH 7.4) and subjected to gentle shaking for three 5-minute washes to eliminate any residual unbound components. DAPI staining solution was applied to the sections, followed by incubation at room temperature in the dark for 10 minutes to facilitate nuclear counterstaining. Finally, anti-fluorescence quenching mounting medium was applied, and the sections were coverslipped. The fluorescent signals were visualized, and high-resolution images were acquired via a multichannel fluorescence scanner capable of detecting the distinct emission spectra of the employed fluorophores.

### Immunohistochemistry

Formalin-fixed paraffin-embedded (FFPE) tumor tissue sections from patients with lung cancer (LC), colorectal cancer (CRC), breast cancer (BT), and ovarian cancer (OC) were dewaxed with xylene and then washed successively with 100% ethanol, 95% ethanol, 75% ethanol, 60% ethanol, and H_2_O for 2 minutes each to complete the hydration process. The glass slide was subsequently heated at 95 °C with sodium citrate solution for 45 minutes. The mixture was subsequently reduced with 3% hydrogen peroxide for 15 minutes and washed with PBS for 3 minutes, for a total of 3 times. Five percent sheep serum was applied to the cover slip tissue. After being blocked and removed unless otherwise specified, the sections were incubated with anti-control, anti-hCD209, anti-CD8 and anti-CD45 antibodies for 1 hour at room temperature. After incubation with a biotin-conjugated goat anti-mouse/rabbit IgG secondary antibody and streptavidin-HRP, positive signals were visualized with a DAB kit (BD Pharmingen) and counterstained with Harris hematoxylin (Fisher Scientific). The formalin-fixed, paraffin-embedded (FFPE) tissue sections were subjected to immunohistochemical staining and subsequently mounted with neutral resin. The entire histological slide was scanned at 20 × magnification via a high-resolution digital pathology scanner. Afterward, 10--20 random high-power fields (HPFs) were selected across the scanned whole-slide image (WSI). The cell-associated chromogenic reaction products were enumerated within each HPF via ImageJ (Fiji) image analysis software. The mean particle density per HPF was calculated as a semiquantitative proxy for molecular expression levels. For quantitative analysis of lymphocyte subsets, the CD8^+^ T-cell and CD209^+^ cell counts were normalized to the CD45^+^ leukocyte count within the same HPF to determine the percentage relative to the total immune infiltrate. The Pearson correlation coefficient was calculated to assess the colocalization relationship between CD8 and CD209 immunostaining patterns within the TME.

### Bioinformatics analysis

The likelihood of immune-checkpoint-blockade response in lung, ovarian, brain, melanoma, and intestinal cancers patients stratified by CD209 expression was predicted with the Tumor Immune Dysfunction and Exclusion (TIDE) algorithm (https://www.aclbi.com/static/index.html#/immunoassay). Survival analyses comparing high- versus low-CD209 expression across tumor types were performed with data from (https://kmplot.com/analysis/, https://cide.ccr.cancer.gov/)(67).

### Statistics

Statistical significance between two groups was determined by an unpaired two-tailed t test; multiple-group comparisons were performed via one-way ANOVA; the weight change curve was analysed via two-way ANOVA for multiple comparisons; and the survival rate was analysed via the log-rank (Mantel‒Cox) test. P <0.05 was considered to indicate statistical significance. The results are shown as the means, and the error bars represent the standard errors of the means (S.E.M.) of biological or technical replicates, as indicated in the figure legends.

### Data and code availability

The single-cell RNA-sequencing data generated from precancerous tissues and primary cancerous tissues from patients with lung adenocarcinoma have been deposited in the Gene Expression Omnibus (GEO) and assigned the accession number GSE298300. The dataset is currently under private status. During peer review, the raw data can be accessed using the following GEO reviewer link and access token:

Reviewer link: https://www.ncbi.nlm.nih.gov/geo/query/acc.cgi?acc=GSE298300 GEO reviewer access token: cpgpcoqcdvmjluf

The single-cell RNA-sequencing data generated from MC38 subcutaneous tumors have been deposited in the Gene Expression Omnibus (GEO) and assigned the accession number GSE308465. The dataset is currently under private status. During peer review, the raw data can be accessed using the following GEO reviewer link and access token:

Reviewer link: https://www.ncbi.nlm.nih.gov/geo/query/acc.cgi?acc=GSE308465 GEO reviewer access token: wfwzsomurvwnxon

The mass spectrometry proteomics data have been deposited to the ProteomeXchange Consortium via the iProX partner repository with the dataset identifier PXD074206.

Reviewer link: https://www.iprox.cn/page/PSV023.html;?url=1777518362467pkdQ Reviewer access token: 5t4d

## Author contributions

T.P. and J.Q.W. performed the experiments with the assistance of Y.Y.L., M.Y., R.R.H., L.Y.F., B.Z., Z.H.C., H.P.W., G.L.H., P.Y.S., Y.W., X.L. and Y.Y.D. T.T.W. and Z.L. helped to perform the glioma experiments; X.X. helped to obtain the human cancer samples and scored the percentages of CD209-and CD8-expressing cells in the samples. T.P. and C.H.W. designed the experiments and analysed the data; C.H.W. wrote the manuscript and supervised the project with Y.Y.D., Z.L. and X.X.

## Acknowledgments

This investigation was supported by grants from the Key Project of the National Natural Science Foundation of China (82430076, to C.H.W.); the National Science Fund for Distinguished Young Scholars (82225029, to C.H.W.); the Special project of the National Natural Science Foundation of China (82441031, to C.H.W.); the General Program of National Natural Science Foundation of China (Grant No. 82572023 to R.R.H.); the Youth Fund of the National Natural Science Foundation of China (82502806, to M.Y, 82302628, to Y.Y.D., 82301989, to R.R.H., 82301987 to B.Z. and 82402704 to Y.Y.L.); The grants from the Department of Science and Technology of Sichuan Province (2026NSFSC1929 to B.Z.); The grant from Sichuan Science and Technology Program (2025JDRC0037 to Y.Y.D.) and the grant from Sichuan Province Innovative Talent Funding Project for Postdoctoral Fellows (BX202516 to H.P.W.).

## Material availability

All the materials generated in this study are available upon reasonable request to the lead contact.

## Figure legends

**Fig. S1. CD209 is selectively upregulated in tumor-infiltrating immunosuppressive myeloid cells and associated with poor clinical response to PD-1 blockade and elevated TIDE scores**

**(A-C)** Clinical validation of CD209 expression in relation to anti–PD-1 therapy response. Tumor samples from patients receiving anti–PD-1 therapy (n = 47) were analyzed by immunohistochemistry.

**(A)** Representative CD209 staining in tumors from anti–PD-1–sensitive and anti–PD-1–resistant patients. Scale bars 200 μm, as indicated.

**(B)** Quantification of CD209⁺ cells among CD45⁺ immune cells in sensitive versus resistant groups.

**(C)** Receiver operating characteristic (ROC) curve evaluating the predictive performance of CD209 expression for anti–PD-1 response.

**(D)** UMAP visualization of single-cell RNA sequencing (scRNA-seq) data from two paired human lung adenocarcinoma and adjacent normal tissues, showing major cell populations.

**(E)** Feature plot displaying CD209 expression across all cell clusters.

**(F)** Comparison of CD209 expression in myeloid subsets (M1, M2 macrophages, monocytes, and dendritic cells) between tumor and adjacent normal tissues.

**(G)** Single-cell RNA sequencing analysis of four paired tumor (LT) and adjacent normal (LN) tissues from lung adenocarcinoma patients (n = 4 pairs). UMAP visualization showing major cell populations, with myeloid cell clusters highlighted.

**(H)** Feature plot showing CD209 expression across cell clusters, indicating preferential expression in myeloid populations.

**(I)** Quantification of CD209 expression in indicated myeloid subsets (M1, M2 macrophages, monocytes, and dendritic cells) comparing tumor (LT) and adjacent normal tissues (LN).

**(J–N)** Association between CD209 expression and immunotherapy response across multiple cancer types. Patients were stratified into CD209-high and CD209-low groups, and tumor immune dysfunction and exclusion (TIDE) scores were compared in colorectal cancer (G), prostate adenocarcinoma (H), breast invasive carcinoma (I), head and neck squamous cell carcinoma (HNSC, J), and uterine corpus endometrial carcinoma (UCEC, K). Statistical significance is indicated.

Statistical significance was determined using two-tailed unpaired t tests (B). *P < 0.05; **P < 0.01; ***P < 0.005; ****P < 0.001. Data in (A-C) are pooled from two independent experiments.

**Fig. S2. Screen for CD209 Abs that rescue human and mouse T-cell activation *in vitro* and exert antitumor activity *in vivo***

**(A)** Binding of monoclonal antibodies to human CD209. 293T cells stably expressing membrane-bound hCD209 were incubated with control IgG or individual anti-hCD209 monoclonal antibodies, followed by FITC-conjugated anti-mouse IgG secondary antibody. Antibody binding was assessed by flow cytometry.

**(B–D)** Functional screening of anti-CD209 monoclonal antibodies in T-cell activation assays. Mouse CD4⁺ T cells (B), mouse CD8⁺ T cells (C), and Jurkat T cells (D) were stimulated with plate-bound αCD3 and αCD28 in the presence of Fc control, hCD209-Fc, or hCD209-Fc supplemented with supernatants from individual hybridoma clones. IL-2 production was measured by ELISA.

**(E)** B16 melanoma cells were intravenously injected into hCD209 knock-in (KI) mice, followed by treatment with control IgG or anti-hCD209 monoclonal antibody. Representative images of metastatic tumor nodules in lungs and livers and their quantification are shown (n = 12 per group).

**(F, G)** GL261-Luc cells were intracranially implanted into hCD209-KI mice. Mice were treated with control antibody, anti-hCD209 monoclonal antibody, temozolomide (TMZ: 100 mg kg⁻¹), or combination therapy following focused ultrasound. (F) Representative bioluminescence imaging at the indicated time points after tumor implantation. (G) Kaplan–Meier survival analysis of treated mice.

Statistical significance was determined using two-tailed unpaired t tests (B–E) and log-rank (Mantel–Cox) test (G). Data are presented as mean ± SEM. *P < 0.05; **P < 0.01; ***P < 0.005; ****P < 0.001. Data in (A-G) are pooled from two independent experiments.

**Fig. S3. Blocking CD209 promotes an antitumour immune response in the TME**

**(A–C)** Single-cell RNA sequencing analysis of tumor-infiltrating immune cells following CD209 blockade.

**(A)** Heatmap showing scaled expression of representative marker genes across major cell populations. Cell clusters are ordered according to the UMAP clustering shown in Figure 3A.\

**(B)** Heatmap of gene expression profiles across T-cell subsets, aligned with the UMAP clustering shown in Figure 3B.

**(C)** Heatmap showing transcriptional programs of tumor-associated macrophage (TAM) subsets, aligned with the UMAP clustering shown in Figure 3C.

**(D–F)** Quantification of immune cell composition derived from the single-cell datasets shown in Figure 3.

**(G)** Flow cytometry gating strategy for identifying tumor-infiltrating CD4⁺ and CD8⁺ T cells and assessing IFN-γ production.

**(H–I)** Frequency of CD4⁺ and CD8⁺ T cells among tumor-infiltrating CD3⁺ T cells in B16 (H) and ID8 (I) tumor models following CD209 blockade.

**(J–K)** IFN-γ production by tumor-infiltrating CD4⁺ and CD8⁺ T cells in B16 (J) and ID8 (K) tumor models.

**(L–M)** IFN-γ levels measured by ELISA in serum and tumor supernatants from MC38 (L) and LLC (M) tumor-bearing mice treated with control or anti-CD209 antibody.

**(N)** IFN-γ production in serum from B16 and ID8 tumor-bearing mice treated with control or anti-CD209 antibody.

Statistical significance was determined using two-tailed unpaired t tests (D-F, H-N). Data are presented as mean ± SEM. *P < 0.05; **P < 0.01; ***P < 0.005; ****P < 0.001. Data in (A-F) are pooled from scRNA-seq experiment with n = 3 tumors per group. Data in (G-N) are pooled from two independent experiments.

**Fig. S4. Blocking CD209 reduces MDSCs but does not alter monocytes or DCs in the tumor microenvironment**

**(A)** Flow cytometry analysis of CD209 expression in splenic and tumor-infiltrating immune cells. Representative gating strategy showing CD209⁺ cells and their distribution among macrophages (F4/80⁺CD11b⁺), monocytes (Ly6C⁺CD11b⁺), and dendritic cells (CD11c⁺MHC-II⁺).

**(B)** Quantification of CD209 expression in total CD45⁺ immune cells and in specific myeloid subsets, including macrophages, monocytes, and dendritic cells, in spleen and tumor tissues. Each dot represents an individual mouse.

**(C)** Flow cytometry gating strategy for identification of myeloid-derived suppressor cells (MDSCs; CD45⁺CD11b⁺Gr-1⁺) and monocytes (CD45⁺CD11b⁺Ly6C⁺) from tumor tissues.\

**(D–E)** Quantification of MDSCs (D) and monocytes (E) among CD45⁺ cells in MC38, LLC, and ID8 tumor models following treatment with control or anti-CD209 monoclonal antibody.

**(F)** Flow cytometry gating strategy for dendritic cells (DCs; CD45⁺CD11c⁺MHC-II⁺) from tumor tissues.

**(G)** Quantification of dendritic cells among CD45⁺ immune cells in MC38, LLC, and ID8 tumor models following CD209 blockade.

Statistical significance was determined using two-tailed unpaired t tests (B, D-E, G). Data are presented as mean ± SEM. ns, not significant; *P < 0.05; **P < 0.01; ***P < 0.005; ****P < 0.001. Data in (A-G) are pooled from two independent experiments.

**Fig. S5. Anti-CD209 antibodies inhibit CD209 function by competitively disrupting ICAM-2 engagement**

**(A)** Immunoblot analysis of ICAM3 expression in control (Vector) and ICAM3-knockout (KO-1, KO-2, KO-3) cells.

**(B)** T cells were stimulated with plate-bound αCD3 and αCD28 in the presence or absence of hCD209-Fc, using control or ICAM3-deficient target cells. IL-2 production was measured by ELISA.

**(C)** Co-immunoprecipitation assays were performed using hCD209-His and mICAM2-Fc in the presence of control or anti-CD209 monoclonal antibodies. Immunoblotting with anti-Fc and anti-His antibodies demonstrates the interaction between CD209 and ICAM2.

**(D–E)** Representative flow cytometry gating strategy and quantification of ICAM2 expression on murine (D) and human (E) CD4⁺ and CD8⁺ T cells.

**(F)** Predicted binding modes and key interacting residues are shown, highlighting potential contact sites between CD209 and ICAM2.

**(G)** Summary of predicted binding residues of CD209 involved in interaction with ICAM2 or anti-CD209 monoclonal antibody, based on structural modeling (AlphaFold3).

**(H)** Binding affinity of wild-type (WT) and mutant CD209 proteins (Mut15) to anti-hCD209 was assessed.

**(I)** Schematic illustration of the T–APC conjugation assay.

Statistical significance was determined using two-tailed unpaired t tests (B). Data are presented as mean ± SEM. *: P<0.05; **: P<0.01; ***: P<0.005; ****: P<0.001. Data in (A-C) are pooled from two independent experiments.

**Figure S6. CD209 blockade mediates antitumor immunity independently of Fc effector functions**

**(A)** Coomassie blue staining of the purified humanized anti-CD209 neutralizing antibody.

**(B)** Surface plasmon resonance (SPR) analysis of the binding affinity between the humanized anti-hCD209 antibody and human CD209 antigen (KD = 2.7 × 10⁻⁹ M).

**(C–E)** MC38 tumors were subcutaneously implanted into hCD209 knock-in (KI) mice, followed by treatment with control hIgG4 or anti-hCD209 hIgG4. Tumor growth curves (C) and tumor weights (D) are shown (n = 12 per group). (E) Flow cytometry analysis of IFN-γ⁺ CD4⁺ and CD8⁺ T cells in the tumor microenvironment.

**(F–H)** Antitumor efficacy of humanized anti-hCD209 antibody in the LLC tumor model. Tumor growth (F) and tumor weights (G) are shown (n = 14 per group). (H) Frequency of IFN-γ⁺ CD4⁺ and CD8⁺ T cells in tumors following treatment.

**(I–L)** Mice were intraperitoneally injected with ID8 cells and treated with control or humanized anti-hCD209 antibody. Representative images of tumor-bearing abdomens (I), abdominal width (J), and ascites volume (K) are shown (n = 12 per group). (L) IFN-γ production by tumor-infiltrating CD4⁺ and CD8⁺ T cells.

**(M–O)** GL261-Luc cells were intracranially implanted into hCD209-KI mice, followed by treatment after focused ultrasound. (M) Representative bioluminescence imaging at indicated time points. (N) Kaplan–Meier survival analysis. (O) Body weight changes during treatment (n = 12 per group).

**(P–Q)** Flow cytometry gating strategies. (P) Identification of CD4⁺, CD8⁺, and IFN-γ⁺ CD8⁺ T cells in the tumor microenvironment. (Q) Identification of tumor-infiltrating myeloid cell subsets, including dendritic cells, macrophages, and monocytes.

Statistical significance was determined using two-way ANOVA (C, F, O), two-tailed unpaired t tests (D–E, G–H, J–L), and log-rank (Mantel–Cox) test (N). Data are presented as mean ± SEM. *: P<0.05; **: P<0.01; ***: P<0.005; ****: P<0.001. Data in (C-Q) are pooled from two independent experiments.

**Figure S7. CD209⁺ myeloid cells inhibit CD8⁺ T-cell infiltration and are associated with poor prognosis across multiple human cancers**

**(A)** Flow cytometry analysis of CD209 expression in tumor-infiltrating immune cells. Representative gating strategy showing CD209⁺ cells within CD45⁺ populations and their distribution across macrophages (CD11b⁺CD68⁺), monocytes (CD14⁺), and dendritic cells (HLA-DR⁺).

**(B)** Quantification of CD209 expression across myeloid subsets.

**(C–F)** Immunohistochemical staining of CD209, CD8, and CD45 in human tumor specimens. Representative images from lung adenocarcinoma (LUAD, C), glioma (D), colorectal cancer (CRC, E), and ovarian cancer (F) patients are shown (P1–P2). Scale bars, as indicated.

**(G–J)** Kaplan–Meier analysis of overall survival in the indicated cancer cohorts. Patients were stratified into CD209-high and CD209-low groups based on the upper and lower quartiles of CD209 expression, respectively. Survival differences were evaluated using the log-rank test.

**(K–N)** Cox proportional hazards analysis of overall survival using CD209 expression as a continuous variable without predefined cutoff. Hazard ratios (HRs), 95% confidence intervals (CIs), and P values were calculated using two-sided Wald tests.

All error bars represent the S.E.M. of technical replicates. The data are representative of two independent experiments (A-F).

