## Supplementary Figures for "Myeloid DC-SIGN drives tumor immune evasion and resistance to PD-1 blockade through ICAM-2–ERM–mediated T-cell mechanosuppression"

**Fig. S1: CD209 is selectively upregulated in tumor-infiltrating immunosuppressive myeloid cells and associated with poor clinical response to PD-1 blockade and elevated TIDE scores**

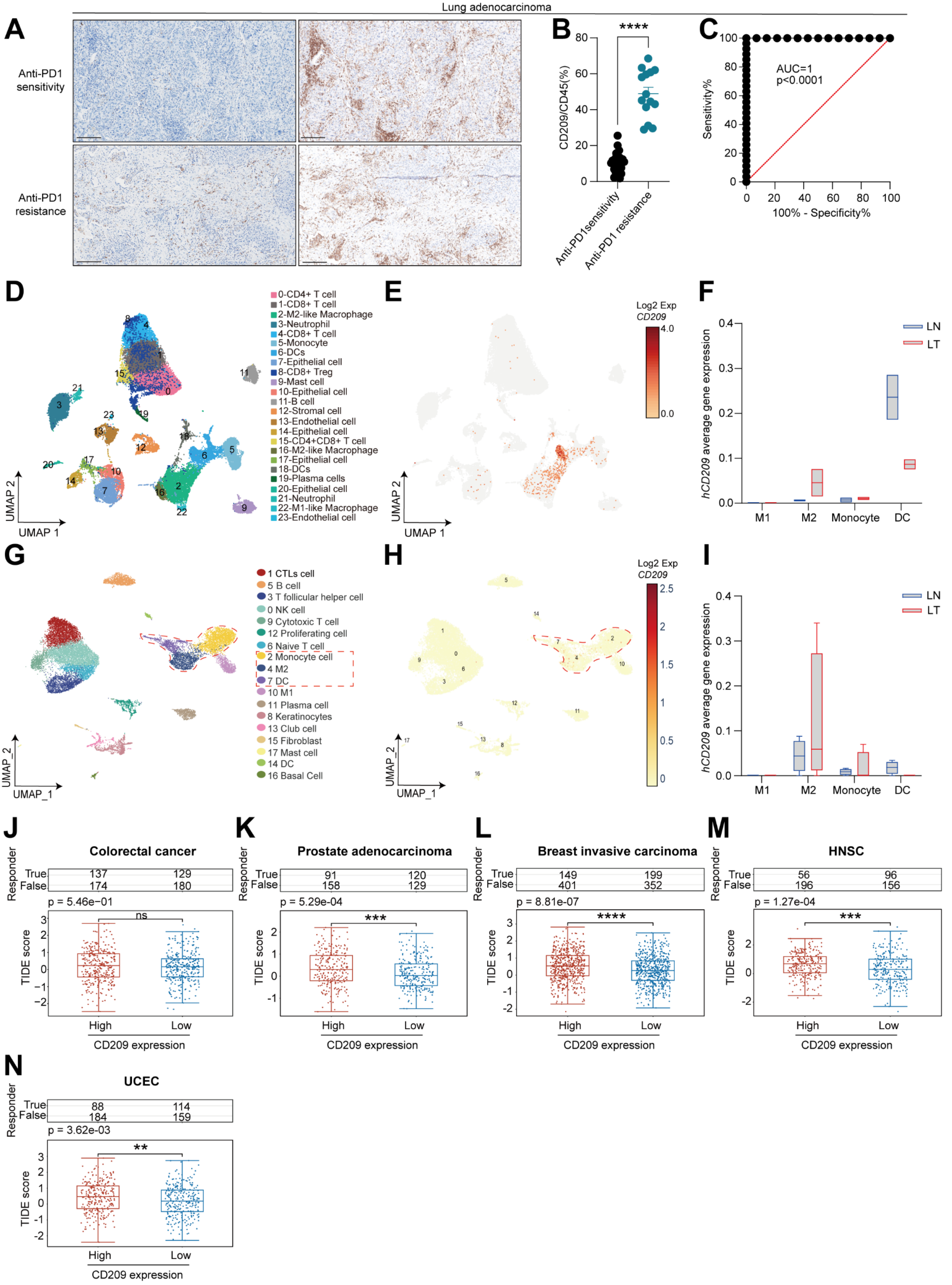

**Fig. S2: Screen for CD209 Abs that rescue human and mouse T-cell activation *in vitro* and exert anti-tumor activity *in vivo***

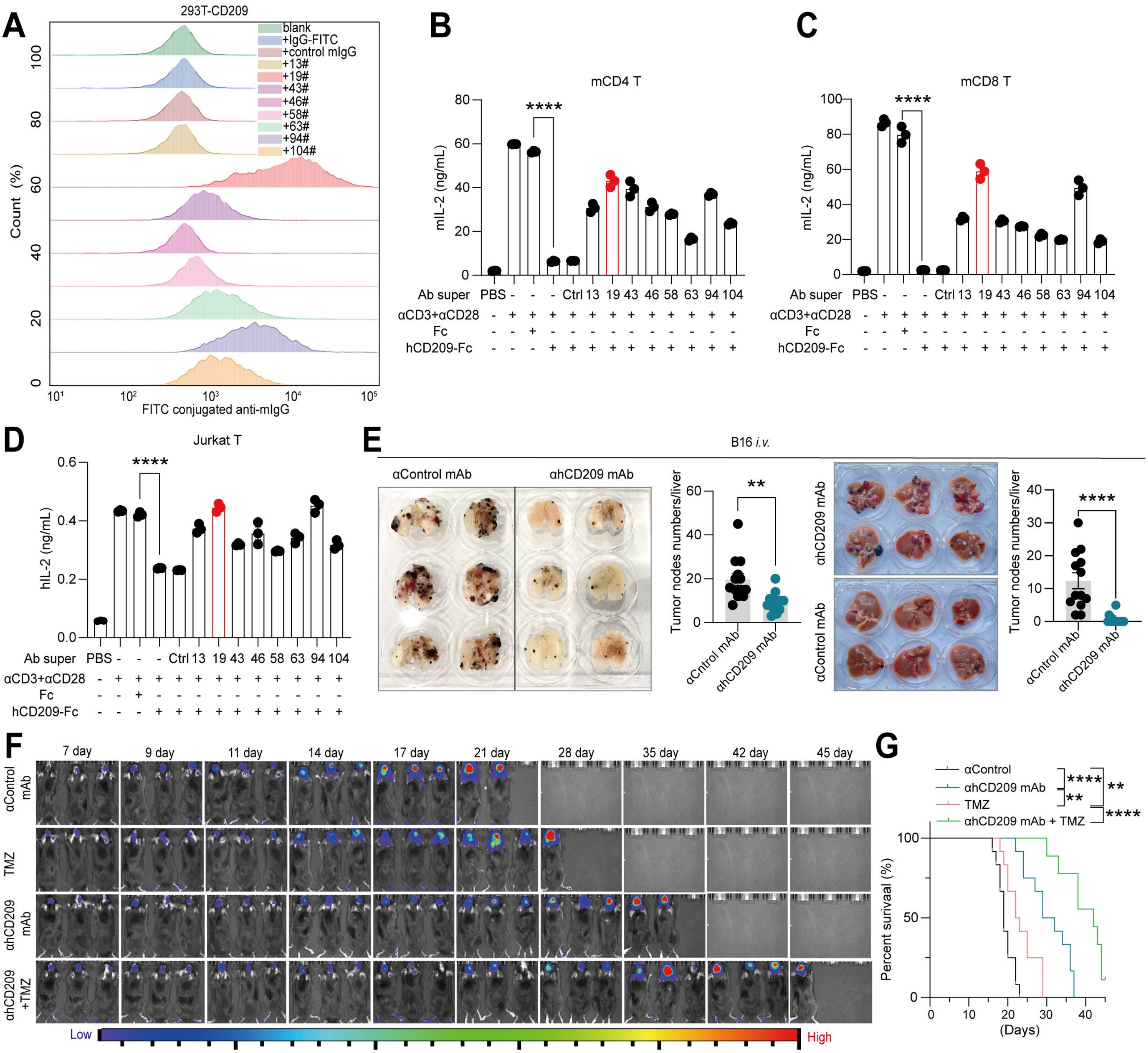

### Fig. S3: Blocking CD209 promotes an antitumour immune response in the TME

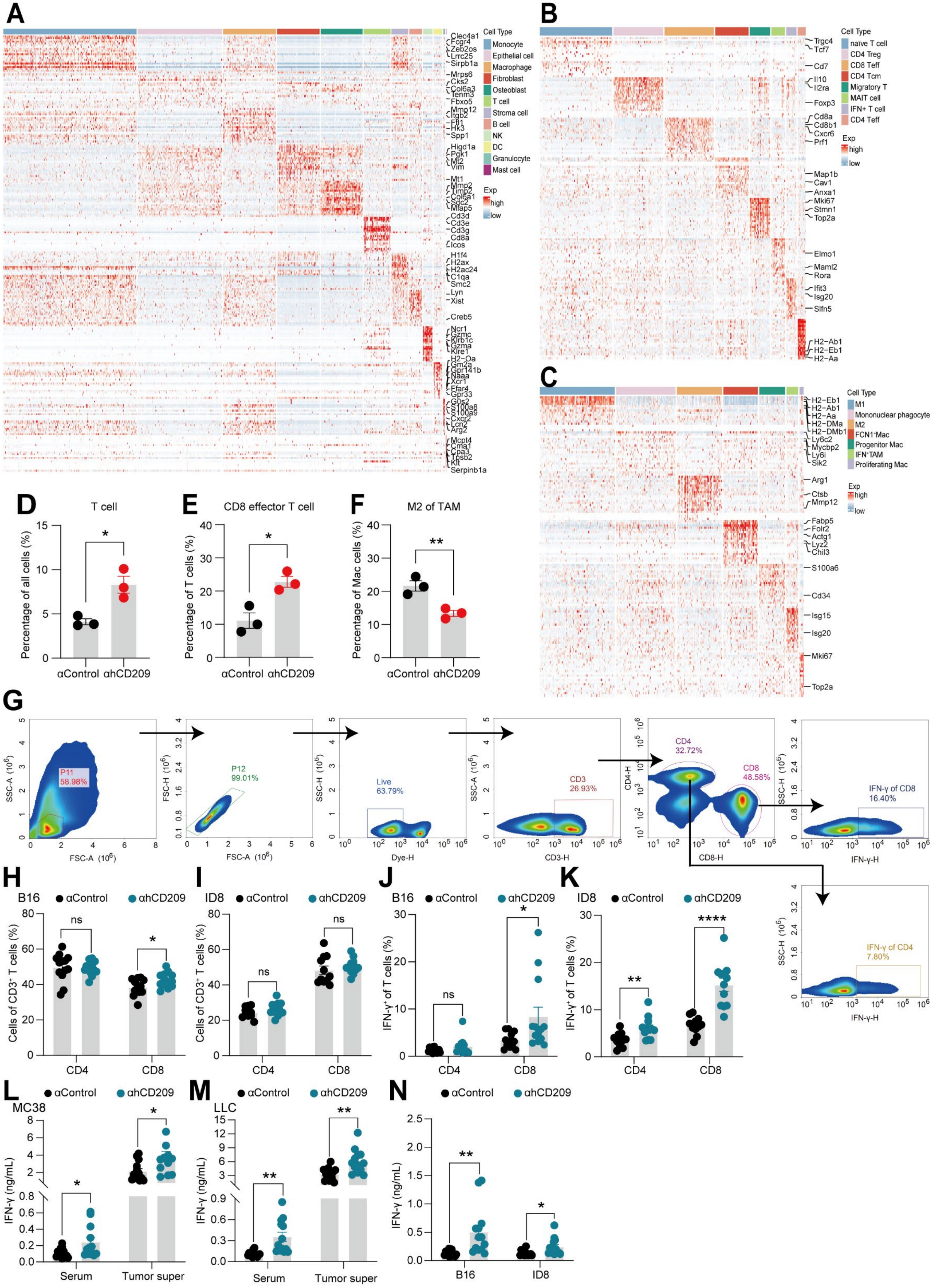

**Fig. S4: Blocking CD209 reduces MDSCs but not change monocytes and DCs in the tumor microenvironment**

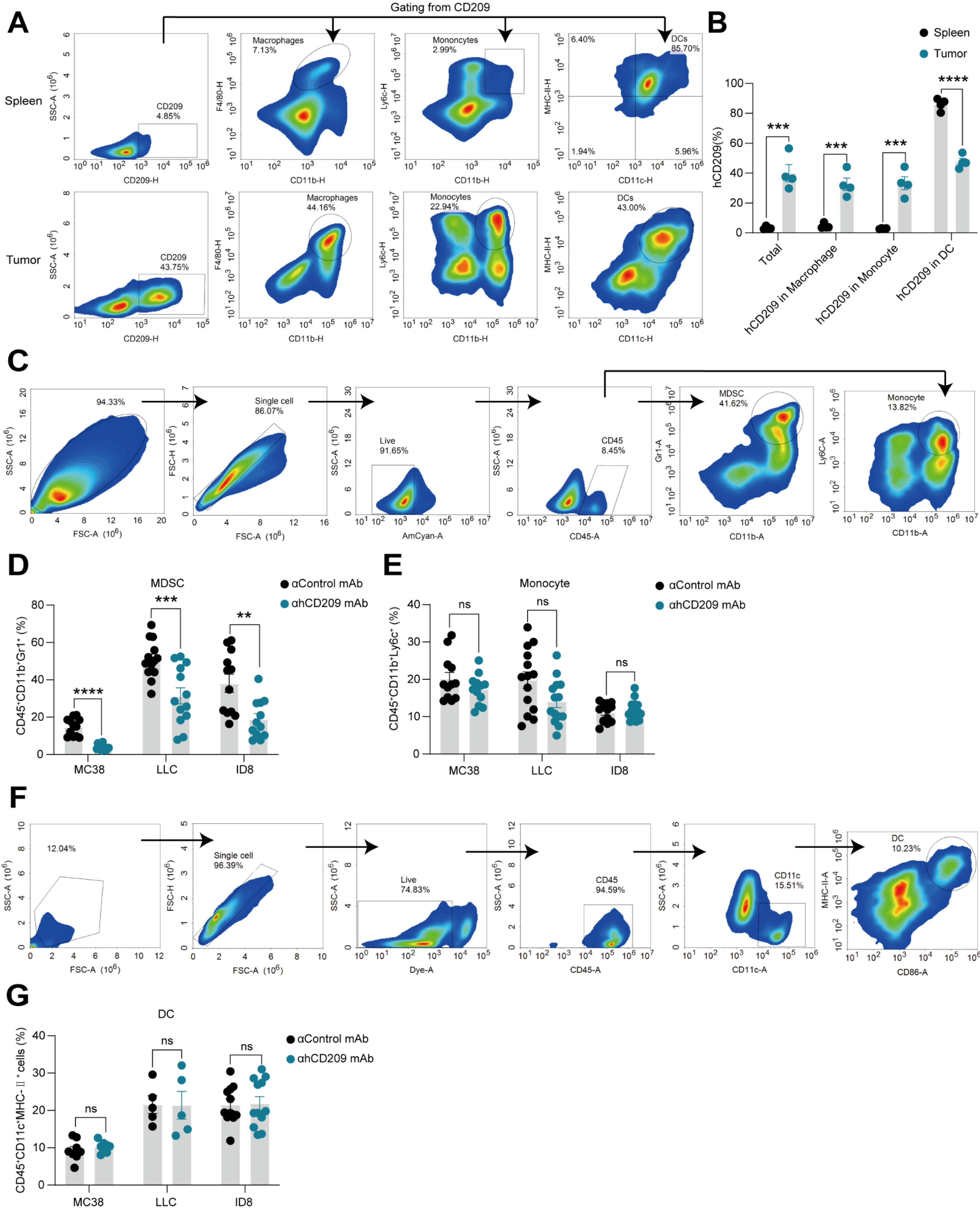

**Fig. S5: Anti-CD209 antibodies inhibit CD209 function by competitively disrupting ICAM-2 engagement**

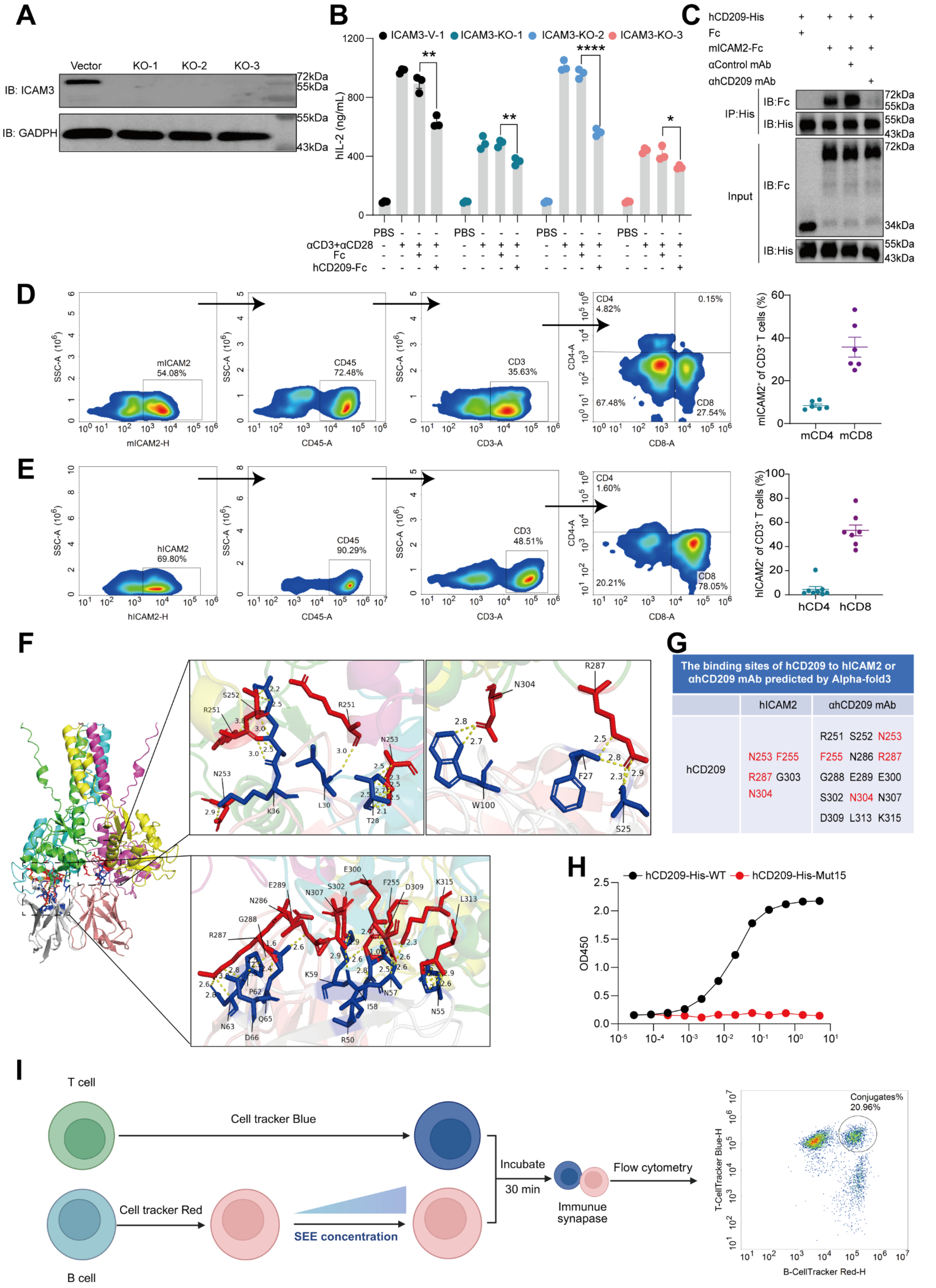


G

| The binding sites of hCD209 to hICAM2 or αhCD209 mAb predicted by Alpha-fold3 | | |
|---|---|---|
| | hICAM2 | αhCD209 mAb |
| hCD209 | N253 F255 R287 G303 N304 | R251 S252 N253 F255 N286 R287 G288 E289 E300 S302 N304 N307 D309 L313 K315 |

**Fig. S6: CD209 blockade mediates antitumor immunity independently of Fc effector functions**

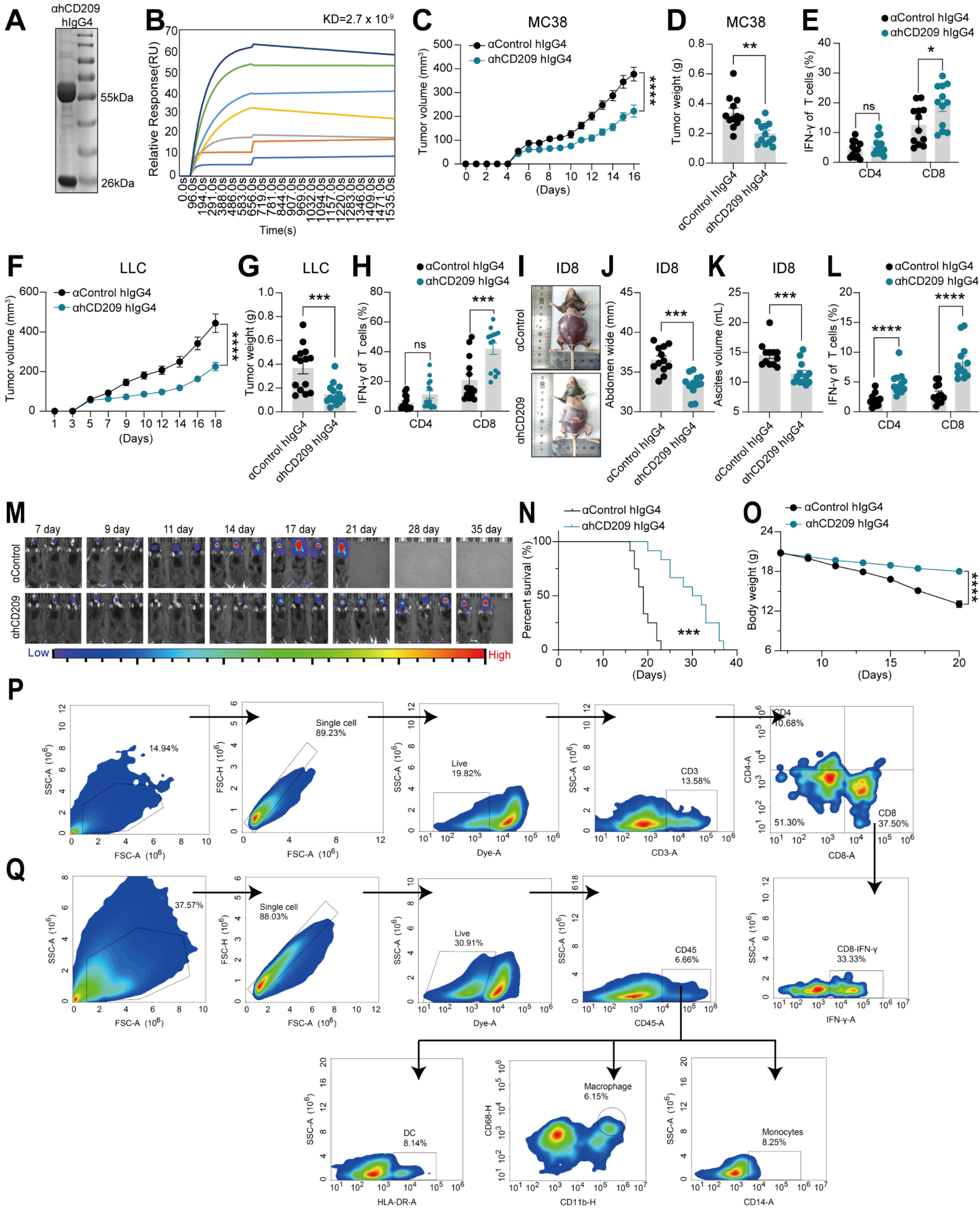

**Fig. S7: CD209$^+$ myeloid cells inhibit CD8$^+$ T-cell infiltration and are associated with poor prognosis across multiple human cancers**

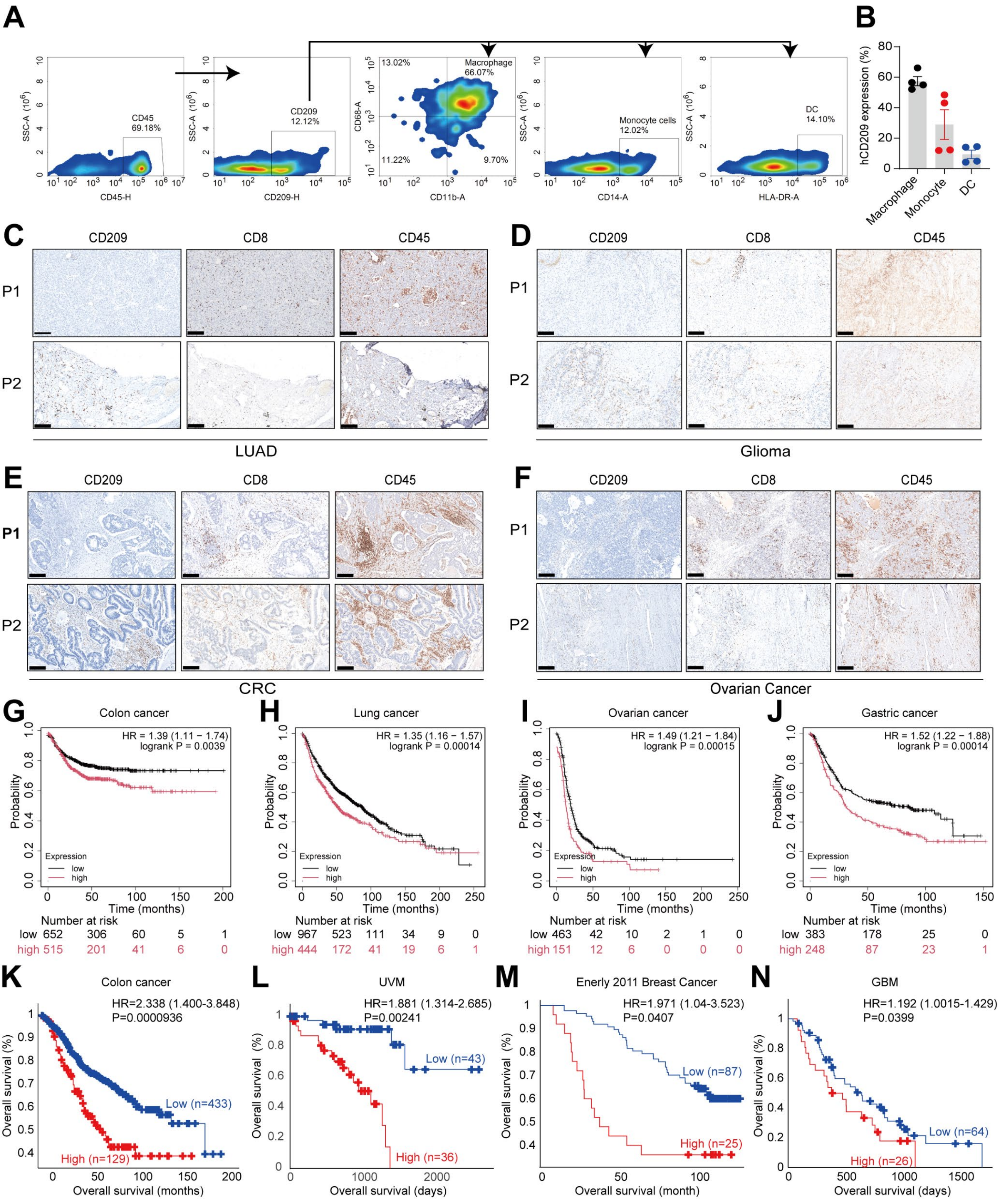

**Table. S1: Knockout gRNA primers for hICAM2 and hICAM3.**

| Target gene | gRNA | Sequence |
|---|---|---|
| hICAM2 | gRNA1 | 5'-CACCGAATACCTTCTCATCCGATCC-3' |
| | | 5'-AAACGGATCGGATGAGAAGGTATTC-3' |
| | gRNA2 | 5'-CACCGTTTTCCAGGATCGGATGAGA-3' |
| | | 5'-AAACTCTCATCCGATCCTGGAAAAC-3' |
| | gRNA3 | 5'-CACCGAGGTATTCGAGGTACACGTG-3' |
| | | 5'-AAACCACGTGTACCTCGAATACCTC-3' |
| hICAM3 | gRNA1 | 5'-CACCGTCATGTTCCTGAAGGTGTCC-3' |
| | | 5'-AAACGGACACCTTCAGGAACATGAC-3' |
| | gRNA2 | 5'-CACCGCATGTTCCTGAAGGTGTCCA-3' |
| | | 5'-AAACTGGACACCTTCAGGAACATGC-3' |
| | gRNA3 | 5'-CACCGTCCTGAAGGTGTCCAGGGGC-3' |
| | | 5'-AAACGCCCCTGGACACCTTCAGGAC-3' |
